# KMT2C and KMT2D regulate skeletal development through stage-specific epigenetic control of chondrogenesis

**DOI:** 10.1101/2025.07.02.662767

**Authors:** Gabrielle Quickstad, Dimitrios V. Bikas, Sara Vardabasso, Karl B. Shpargel

## Abstract

Many craniofacial disorders linked to mutations in enhancer-associated chromatin-modifying enzymes, including Kabuki syndrome (KS), present with a wide range of skeletal abnormalities. KS is a craniofacial development disorder characterized by mutations in KMT2D, a histone H3 lysine 4 (H3K4) methyltransferase. The KMT2D cellular origins and molecular pathways leading to skeletal deficits in KS are not well characterized. We previously demonstrated that KMT2D deletion in mouse cranial neural crest cells (cNCCs), the multipotent stem cells that contribute to many craniofacial structures, specifically impairs chondrocyte hypertrophic differentiation. Although we observed defects in cNCC endochondral ossification, broader skeletal function of KMT2D is largely unknown and its functional redundancy in skeletal development with its paralog, KMT2C, is understudied. We now find that KMT2D mutation within chondrocyte lineages results in postnatal growth defects and impaired skeletal bone development. We demonstrate redundancy of KMT2C in regulating endochondral ossification as mice lacking both methylases within chondrocyte lineages demonstrate severe deficits in bone formation. These methylases regulate change in chondrocyte state as hypertrophic differentiation is impaired with loss of both KMT2C and KMT2D. When both methylases are deleted in stem cell lineages, KMT2C/KMT2D double mutant cNCCs are unable to properly induce chondrocyte factors downstream of SOX9 activation. At the molecular level, there is a loss of H3K4me1 at de novo chondrocyte enhancers essential for skeletal development and an associated reduction in the efficiency of acetylation with KMT2C/D deficiency, suggesting partial dependency on KMT2C/D presence for appropriate deposition of H3K27ac. We propose a role for KMT2C and KMT2D in facilitating proper activation of genes downstream of SOX9 and RUNX2 transcription factors. Thus, we delineate a regulatory cascade orchestrated by chromatin-level modifications wherein KMT2C/D activity ensures commitment of chondrocyte lineages via enhancer priming. This work not only expands the functional repertoire of COMPASS family members but also defines a chromatin-based mechanism for vertebrate skeletal development, offering insights into the epigenetic basis of skeletal dysplasia and related congenital disorders.

## Introduction

Kabuki Syndrome (KS) is a craniofacial development disorder whereby patients display facial gestalt including a broad depressed nasal tip, high arched eyebrows, and a high-arched or cleft palate (Niikawa et al., 1988). The disorder can also include the development of multi-organ deficiencies and skeletal abnormalities. A summary of 13 different clinical reports indicated that 55% of patients presented with short stature and 88% presented with skeletal abnormalities (Matsumoto & Niikawa, 2003). Rib and vertebral anomalies have been observed, including sagittal cleft, butterfly vertebrae, and hemivertebrae (Matsumoto & Niikawa, 2003). Patients can also present with cleft hand, brachydactyly and scoliosis, as well as joint hypermobility and dislocations (Kuroki et al. 1981; Niikawa et al. 1988, 1981; Schrander-Stumpel et al. 2005; Adam, Hudgins, and Hannibal 1993; Cocciadiferro et al. 2018; Hannibal et al. 2011)(Huh et al., 2011). Together, these patient reports confirm that skeletal abnormalities are a defining feature of KS.

KS is associated with mutations in two distinct histone-modifying enzymes, KMT2D (MLL4) and UTX (KDM6A), with poorly characterized function (Shpargel & Quickstad, 2023). Causative heterozygous mutations in KMT2D account for 39-74% of patient cases (KS1) while mutations in UTX (KS2) are less common (3-6%) (Banka et al., 2015; Nina Bögershausen et al., 2016; Cocciadiferro et al., 2018; Hannibal et al., 2011; Lederer et al., 2012; Makrythanasis et al., 2013; Micale et al., 2011; Miyake et al., 2013; S. B. Ng et al., 2010). KMT2D is a histone H3 lysine 4 methylase which, along with its paralog KMT2C (MLL3), is responsible for depositing H3K4me1, a marker enriched at primed enhancers (Heintzman et al., 2007; Herz et al., 2012; Hu et al., 2013; Lai et al., 2017; Rada-Iglesias et al., 2011; L.-H. Wang et al., 2021). Although KMT2C mutations have not been described in KS, they are implicated in the intellectual disorder Kleefstra syndrome 2 (KLEFS2), which also presents with craniofacial components and short stature (Koemans et al., 2017; Rots et al., 2024; Siano et al., 2022). UTX, a histone H3 lysine 27 demethylase, functions in a complex with KMT2C and KMT2D and is important for the removal of H3K27me3 from poised enhancer regions so the CREBBP/EP300 complex can acetylate this residue (S.-P. Wang et al., 2017). Although other H3K4 methylases comprise the catalytic subunit of COMPASS complexes, UTX only associates with those containing KMT2C or KMT2D (Cho et al., 2007).

Craniofacial disorders often involve mutations in chromatin complexes that participate in COMPASS co-regulation at enhancers. For example, craniofacial disorders such as CdLS, Rubinstein-Taybi, and CHARGE are directly linked to mutations in epigenetic modifiers (NIPBL, CREBBP/EP300, and CHD7, respectively) (Mannini et al., 2013; Roelfsema et al., 2005) (Shangguan & Chen, 2022)(Ufartes et al., 2021)(Negri et al., 2019). Despite having distinct syndromic features and characteristic facial morphology, patients frequently display overlapping traits such as impaired growth and short stature, proposing epigenetic mechanisms for broader disruption of skeletal development. Accordingly, KMT2D mutations have been implicated in other craniofacial disorders.

Some individuals with Cornelia de Lange (CdLS)-like or CHARGE syndrome-like features carry KMT2D mutations (Kaur et al., 2024; Moccia et al., 2018; Sakata et al., 2017; Shangguan & Chen, 2022). These shared phenotypic qualities not only point to epigenetic convergence due to similar mis-regulation of transcription, but also emphasize the importance of KMT2D in craniofacial and skeletal development.

As defects in bone development are considered to be pathological causes of short stature, KMT2D’s role in chondrocyte differentiation could explain KS skeletal pathologies (Garofalo et al., 1999; Warman et al., 2011). A prior study found that short stature was observed in 70% of patients positive for a mutation in *Kmt2d* compared to only 25% of patients carrying other mutations (N Bögershausen & Wollnik, 2013). Most bones develop through endochondral ossification, a process that begins with the condensation of mesenchymal stem cells into chondrocytes, the cell type found in cartilage (Hall & Miyake, 2000). These cells enter a terminal differentiation stage called hypertrophy, forming a scaffold for the primary and secondary centers of ossification to form (Kronenberg, 2003; Mackie et al., 2008). Zebrafish models of KS have shown that deletion of *Kmt2d* is associated with abnormal chondrocyte differentiation (Shangguan et al., 2024); however, mammalian chondrocyte-specific function requires further exploration. Bone biology has been explored in heterozygous *Kmt2d* gene trap mice as loss of KMT2D resulted in expanded growth plates resulting from excessive differentiation and increased zones of proliferative and hypertrophic chondrocytes. Additional *in vitro* data indicated excessive chondrocyte differentiation in the absence of KMT2D (Fahrner et al., 2019).

Previously, we used a mouse model with a neural crest cell-specific deletion of *Kmt2d* to model human KS facial pathology (Shpargel et al., 2020). Cranial neural crest stem cells (cNCCs) are the multipotent stem cells responsible for the development of all anterior facial bone and cartilage. cNCC-derived chondrocytes in the cranial base undergo endochondral ossification similar to the axial and appendicular skeleton, with the exception that growth is bidirectional from synchondroses as opposed to unidirectional growth plates (McBratney-Owen et al., 2008; Szabo-Rogers et al., 2010). In contrast to other studies, we found that cNCC-specific deletion of *Kmt2d* did not alter chondrocyte distributions in resting and proliferative zones within cranial base synchondroses (Shpargel et al., 2020). Rather, KMT2D mutant chondrocytes fail to reach the terminal hypertrophic stage necessary for bone formation, suggesting that KMT2D is specifically involved in this crucial transition stage in chondrocyte differentiation through unknown mechanisms. These conflicting data highlight the need for relevant mammalian *in vivo* and *in vitro* chondrocyte models and to understand KMT2D molecular and cellular function in skeletal development. Chondrocyte terminal differentiation is necessary for the proper development of bone by promoting vascularization, initiating extracellular matrix mineralization, and indirectly and directly allowing for the replacement of cartilage with bone (Long et al., 2022; Sun & Beier, 2014; Yang et al., 2014). Thus, we hypothesize that KMT2D and KMT2C play a broader role in skeletal development during cartilage differentiation, explaining the skeletal pathologies seen in KS and KLEFS2 patients.

In the present study, we utilized a chondrogenic differentiation model *in vitro* and a transgenic mouse line in which expression of Cre recombinase is under the control of the mouse type II collagen (*Col2a1*) promoter or the mouse *Sox10* promoter to generate animals with chondrocyte-specific or cNCC-specific inactivation respectively of *Kmt2c* and/or *Kmt2d*. Our study identifies KMT2C and KMT2D as important epigenetic regulators in the process of stage-specific expression of chondrogenic and osteogenic factors. Genetic ablation of *Kmt2c/d* disrupts the enhancer landscape and prevents the deposition of H3K4me1 at lineage-specific enhancers during chondrocyte differentiation. These changes correspond with profound defects in mesenchymal stem cell differentiation and skeletal patterning, as evidenced by both *in vivo* phenotypic observations and cellular morphology, as well as *in vitro* differentiation defects. Our findings provide evidence that expression of both KMT2C and KMT2D is necessary for chondrocyte specification and hypertrophic differentiation during bone development.

## Results

### KMT2C and KMT2D have partially redundant function in endochondral ossification

We utilized *Kmt2d* and *Kmt2c* conditional alleles with exons floxed upstream of the catalytic SET domains (Fig. S1). Prior research has established that Cre-induced recombination of these alleles results in an absence of KMT2C or KMT2D proteins (J.-E. Lee et al., 2013; S. Lee et al., 2006; Shpargel et al., 2020). To assess the role of KMT2C and KMT2D in endochondral ossification, we crossed *Kmt2c/d* conditional alleles with a *Col2a1-Cre* transgene to generate chondrocyte-specific knockout mice (Fig. 1a). Using a Cre-activated *Rosa^Tomato^*reporter, we observed near complete expression of the Tomato reporter in tibial chondrocytes accompanied by an absence of H3K4me1 (Fig. 1b, indicating a loss of KMT2C/D methylase activity in Cre-expressing chondrocytes.

**Figure 1:**
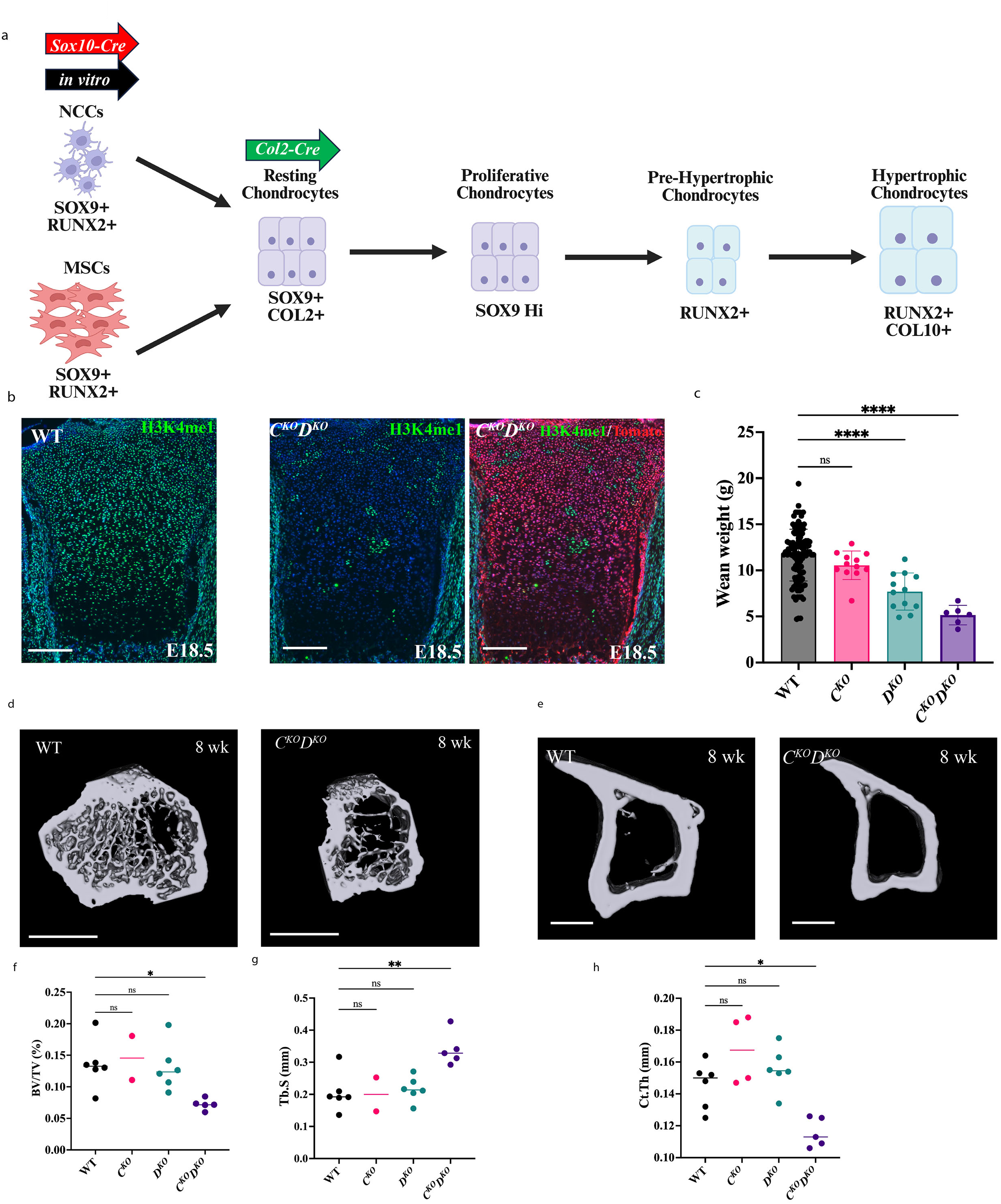
KMT2C and KMT2D play redundant roles in the regulation of bone formation. **a** Model depicting chondrocyte differentiation from both NCC and MSC lineages. *Sox10-Cre* (red arrow), *Col2a1-*Cre (green arrow), and *in vitro* (black arrow) systems are indicated at the stage of differentiation KMT2C/D loss is occurring in each model. Common markers for each stage of chondrocyte differentiation are labeled. **b** Immunofluorescence of coronally sectioned proximal E18.5 tibia in WT and *C^KO^D^KO^* for H3K4me1 and *Rosa^Tomato^*(Tomato) reporter fluorescence revealed near total expression of *Cre* with an associated absence of H3K4me1 in *Rosa^Tomato^* + cells in *C^KO^D^KO^*. Scale bar is 200 µm. **c** Wean weights (P24) in grams (g) of *C^KO^*, *D^KO^*, *C^KO^D^KO^*, and WT (*N*=112, *N*=12, *N*=12, *N*=6, respectively). **d-e** Representative three-dimensional micro-CT reconstructions of 8-week-old WT and *C^KO^D^KO^* proximal tibial region measured for trabecular and cortical analyses. Transverse view shown in the direction of the distal tibial end. Scale bar is 1 mm (d) or 500 µm €. **f-g** Micro-CT analysis of trabecular bone formation (BV/TV, Tb.S) in 8-week-old tibia of WT, *C^KO^*, *D^KO^*, and *C^KO^D^KO^* mice (*N*=6, *N*=5, respectively). **h** Cortical thickness in mm shown for WT, *C^KO^*, *D^KO^*, and *C^KO^D^KO^*. Data shown are mean ± SD. Statistical analyses were performed with Welch ANOVA followed by Dunnett’s T3 multiple comparisons test (for wean weight) or ordinary one-way ANOVA followed by Dunnett’s test (for trabecular analysis). *P<0.05, **P<0.01, ***P<0.001, ****P<0.0001, ns = not significant where specified

Mice carrying chondrocyte-specific double knockout (DKO) of KMT2C and KMT2D (*Kmt2c^fl/fl^ Kmt2d^fl/fl^ Col2a1-Cre* = *C^KO^D^KO^*, see Table S1 for genotype abbreviations) displayed substantial postnatal growth delays, malocclusion, and unsteadiness at weaning (Fig. 1c Table S2). *C^KO^D^KO^* mice were underrepresented at postnatal day 0 (P0) (8 observed vs. 30 expected), suggesting perinatal lethality, whereas genotype frequencies were normal at late embryonic stages (E18.5) (Table S3).

Discoloration of *C^KO^D^KO^* pups at P0 suggests partial penetrance of perinatal lethality due to asphyxia (Fig. S2). Chondrocyte-specific single knockout (SKO) of KMT2D (*Kmt2c^fl/+^ Kmt2d^fl/fl^ Col2a1-Cre* = *D^KO^*) exhibited milder postnatal growth defects and were born at Mendelian ratios, indicating that one allele of *Kmt2c* is sufficient to restore viability and partially compensate for the loss of KMT2D. Notably, KMT2C SKO (*Kmt2c^fl/fl^ Kmt2d^fl/+^ Col2a1-Cre* = *C^KO^*) closely resembled wildtype (WT) Cre-negative mice at weaning by weight and phenotypic observations, providing evidence that KMT2D can sufficiently support growth in the absence of KMT2C.

We collected skeletally juvenile mice at 8 weeks of age for micro-CT analysis to evaluate changes to bone microarchitecture that could be associated with the severe growth defects as *D^KO^*and *C^KO^D^KO^* mice approach adulthood. We performed micro-CT of the proximal tibia to assess trabecular bone volume fraction (BV/TV), number (Tb.N), thickness (Tb.Th), and separation (Tb.S) (Fig. 1d, f-g). Shown are transverse views of representative three-dimensional reconstructions of the proximal tibial region shown in the direction of the distal tibial end. No significant changes were observed between WT and *C^KO^* or *D^KO^*individual mutation; however, *C^KO^D^KO^* mice had a loss of trabeculation. These chondrocyte-specific DKO mice exhibited a significant increase in Tb.S and a significant decrease in relative trabecular bone volume, Tb.N and Tb.Th as compared to WT (Fig. 1f-g). Cortical analyses of the tibial midshaft revealed a similar loss of cortical thickness in *C^KO^D^KO^*(Fig. 1e). These findings suggest that combined loss of KMT2C and KMT2D impairs trabecular and cortical bone formation, likely resulting in the extreme growth deficits observed in *C^KO^D^KO^*mice. Collectively, these data demonstrate that KMT2C and KMT2D can partially compensate for the loss of the other in the context of postnatal bone growth, as losing both methylases results in more severe skeletal defects than losing one alone.

### KMT2C/D-deficient growth plates have deficient columnar organization and hypertrophy

We performed histological analysis of 8-week coronally sectioned proximal tibial growth plates to determine how loss of KMT2C/D impacted chondrocyte organization and cell morphology. Safranin O/fast green staining revealed a thicker growth plate in both *D^KO^* and *C^KO^D^KO^*, with *C^KO^D^KO^* proximal growth plates over 2-fold thicker than WT (Fig. 2a-b). H&E staining indicated a loss of typical chondrocyte columnar organization in both proliferative and hypertrophic zones of *C^KO^D^KO^*(Fig. 2a). A spatial representation of the proliferative columnar distribution of cells indicated a significant loss of linearity in *C^KO^D^KO^* columnar stacking compared to both WT and *D^KO^* (Fig. 2c). The total number of hypertrophic chondrocytes and density of hypertrophic cells were reduced in *D^KO^*and more severely impaired in *C^KO^D^KO^* mice (Fig. 2d-e, Fig. S4a). We also noted zones of safranin O-positive (proteoglycan-expressing) cells expressing chondrocyte markers (RUNX2) embedded within the tibial diaphysis of *C^KO^D^KO^*mice, indicating that KMT2C/D-deficient chondrocytes may be incapable of effectively transitioning to terminal differentiation (Fig. S4b-c). Rather than entering terminal differentiation that leads to programmed chondrocyte death and resorption, these chondrocytes have become embedded within regions of bone formation. Altogether, these findings indicate that KMT2C/D-deficient chondrocytes fail to properly organize tibial growth plates and undergo normal hypertrophic differentiation.

**Figure 2:**
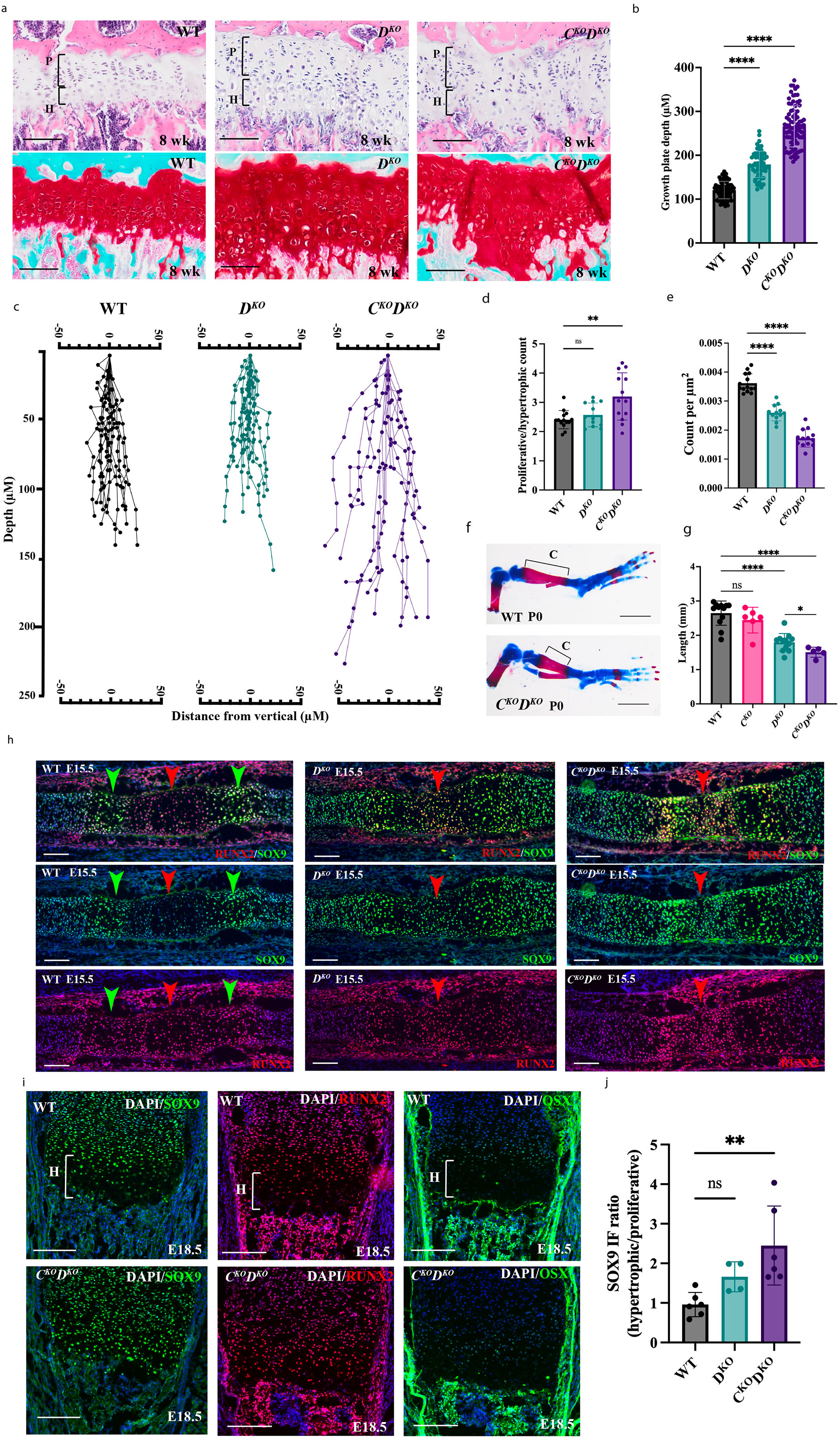
KMT2C/D loss impairs chondrocyte hypertrophy and bone formation. **a** Hematoxylin and eosin (H&E) staining and Safranin O/fast green FCF staining in 8-week-old coronally sectioned proximal tibial growth plates of WT, *D^KO^*, and *C^KO^D^KO^* mice. Proliferative (P) and Hypertrophic (H) regions demarcated by brackets showing increased disorganization in *C^KO^D^KO^* as compared to WT. Scale bar is 100 µm. **b** Eight-week-old proximal tibial growth plate thickness (µM) in WT, *D^KO^*, and *C^KO^D^KO^*(*N*=4, *N*=3, *N*=4, respectively). **c** Quantitation of longitudinal columns of proximal tibial growth plate chondrocytes measuring ten cells down from the vertical origin point in WT, *D^KO^*, and *C^KO^D^KO^* measured in depth (µm) and distance from the vertical origin point (µm). **d-e** The ratio of proliferative to hypertrophic cells and the density of hypertrophic cells in 8-week-old proximal tibial growth plates of WT, *D^KO^*, and *C^KO^D^KO^* (*N*=4, *N*=3, *N*=4, respectively). **f** Alcian blue and alizarin red staining for cartilage and bone of WT and *C^KO^D^KO^*P0 hindlimbs. Calcified region (C) demarcated with brackets. Scale bar is 1 mm. **g** Length (mm) of calcified region of bone in P0 *C^KO^*, *D^KO^*, *C^KO^D^KO^*, and WT, (f) and *D^KO^*as compared to *C^KO^D^KO^* (g)(*N*=12, *N*=6, *N*=12, *N*=5, respectively). **h** Immunofluorescence for SOX9 (green) and RUNX2 (red) in E15.5 tibial diaphysis. Red arrow indicates hypertrophic region, and green arrows indicate proliferative regions. Scale bar is 100 µm. **i** Immunofluorescence for SOX9 (green), RUNX2 (red), or Osterix (OSX green) in E18.5 proximal tibial growth plate of WT and *C^KO^D^KO^*. Hypertrophic region (H) marked with brackets. **j** Quantification of the ratio of SOX9 IF intensity of hypertrophic over proliferative cells in E15.5 WT, *D^KO^*, and *C^KO^D^KO^* (*N*=6, *N*=4, *N*=6). Data shown are mean ± SD. Statistical analyses were performed using ordinary one-way ANOVA followed by Dunnett’s test or unpaired t-test with Welch’s correction for *D^KO^* and *C^KO^D^KO^* comparison. *P<0.05, **P<0.01, ***P<0.001, ****P<0.0001, ns = not significant where specified

### KMT2C and KMT2D regulate bone development through chondrocyte hypertrophy

Given the evidence of disrupted hypertrophic differentiation, a key step in bone growth, in 8-week-old *D^KO^*and *C^KO^D^KO^* mice, we hypothesized that an earlier developmental defect might underlie the impaired skeletal growth. Alizarin red and alcian blue staining of bone and cartilage in newborn pups (P0) displayed a progressive reduction in the length of alizarin red-stained calcified zones within the tibial diaphysis, with the phenotype severity increasing from *C^KO^* to *D^KO^*to *C^KO^D^KO^* relative to WT controls (Fig. 2f-g).

In mice, mesenchymal stem cell (MSC) condensation, chondrocyte specification and proliferation occur between embryonic days E12.5 and E15.5, coinciding with the expression of SOX9, a key transcription factor (TF) required for the initiation and maintenance of proliferating chondrocytes (Bi et al., 1999). By E15.5, chondrocytes begin to enter terminal differentiation, marked by the expression and increased activity of RUNX2, a TF essential for initiating bone formation (Toshihisa Komori, 2010; T Komori et al., 1997). The transition to hypertrophic differentiation is partially regulated by the relative expression and activity of SOX9 and RUNX2 (Ducy et al., 1997; Foster et al., 1994; Inada et al., 1999; I. S. Kim et al., 1999; T Komori et al., 1997; Mundlos et al., 1997; L. J. Ng et al., 1997; Otto et al., 1997; Wright et al., 1995; Yoshida et al., 2004; Zhao et al., 1997; Zhou et al., 2006). Immunofluorescence in *C^KO^D^KO^* E15.5 tibial diaphyses indicate chondrocytes properly specify, expressing SOX9 (Fig. 2h). E15.5 *C^KO^D^KO^*chondrocytes enter the proliferative state, expressing RUNX2 as expected in early pre-hypertrophic chondrocytes; however, these chondrocytes fail to decrease SOX9 protein levels in hypertrophic chondrocytes (Fig. 2h). Quantitation of the ratio of SOX9 protein in hypertrophic chondrocytes relative to proliferative zones indicates a progress enhancement in *D^KO^* and *C^KO^D^KO^*embryos across biological replicates (Fig. 2j). As hypertrophic differentiation progresses, at E18.5, *C^KO^D^KO^*proximal tibial growth plates exhibit a reduction in hypertrophic zone thickness (Fig. 2i). We conclude that KMT2C and KMT2D function redundantly in the regulation of SOX9 expression and chondrocyte hypertrophy during terminal differentiation.

### KMT2C/D promote hypertrophic differentiation within cNCC and mesodermal chondrocytes

The cranial base forms through a combination of mesodermal and cNCC lineages with the presphenoid (PS) and anterior half of the basisphenoid (BS) region deriving from cNCCs (McBratney-Owen et al., 2008). We examined cranial phenotypes in the *Col2a1-Cre* mice to contrast KMT2C and KMT2D function in endochondral ossification within cNCC lineages (hyoid, PS, BS) to those of mesodermal origin (basioccipital = BO, tibia). At birth, *C^KO^D^KO^* hyoid cartilage shows a total loss of alizarin red staining for calcified zones of bone (Fig. S5). The cranial base in *C^KO^D^KO^* displays a dramatic reduction in the areas of the BO, BS, and PS regions of bone with more mild phenotypes in *D^KO^*mice than *C^KO^D^KO^* (Fig. 3a-b). In *C^KO^D^KO^*, there is a complete loss of PS bone at birth. Together, these data indicate an inability of both mesodermal and cNCC derived chondrocytes to appropriately stimulate bone formation.

**Figure 3:**
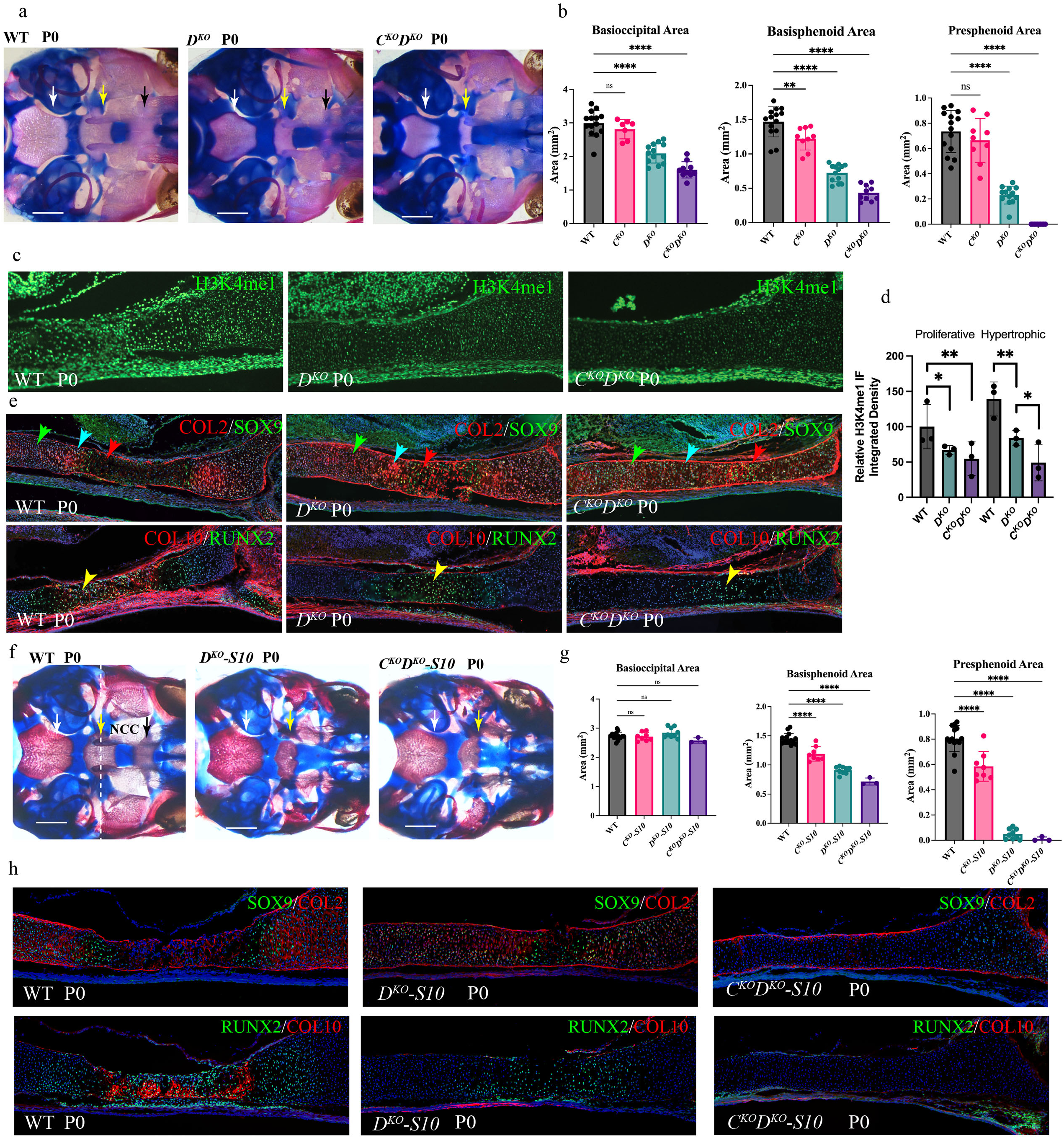
KMT2C/D are required for both chondrocyte specification and maintenance of hypertrophy. **a,f** Alizarin red and alcian blue staining of P0 WT, *D^KO^*, and *C^KO^D^KO^* (a) and WT, *D^KO^-S10*, and *C^KO^D^KO^-S10* (f) with ventral whole mount view of the cranial base. Basioccipital (white arrow), basisphenoid (yellow arrow), and presphenoid (black arrow) are indicated. Scale bar is 1 mm. In part f, NCC-derived bone formation is specified to the right of the white dashed line (the presphenoid region and anterior basisphenoid region). Scale bar is 1 mm. **b,g** Area (mm^2^) of basioccipital, basisphenoid, and presphenoid regions of P0 *Col2a1-Cre* (b: *N*=14, *N*=9, *N*=13, *N*=10, respectively) and *Sox10-Cre* (g: *N*=14, *N*=8, *N*=9, *N*=3, respectively) mice. **c** H3K4me1 immunofluorescence of sagittal section of the presphenoid region of the cranial base in P0 WT, *D^KO^*, and *C^KO^D^KO^* **d** Relative H3K4me1 integrated density in P0 WT, *D^KO^*, and *C^KO^D^KO^* in both proliferative and hypertrophic chondrocytes of the presphenoid region. **e** Immunofluorescence of sagittal section of the presphenoid region in P0 WT, *D^KO^*, and *C^KO^D^KO^* for COL2A1 (red) and SOX9 (green, top panels) and RUNX2 (red) and COL10A1 (green, bottom panels). **h** Immunofluorescence of sagittal section of the presphenoid region in P0 WT, *D^KO^*, and *C^KO^D^KO^* for COL2A1 (red) and SOX9 (green, top panels) and RUNX2 (green) and COL10A1 (red, bottom panels). Data shown are mean ± SD. Statistical analyses were performed using ordinary one-way ANOVA followed by Dunnett’s test or unpaired t-test with Welch’s correction for *D^KO^* and *C^KO^D^KO^*comparison. *P<0.05, **P<0.01, ***P<0.001, ****P<0.0001, ns = not significant where specified

Meckel’s cartilage, a cNCC derived developmental cartilaginous structure forming along the mandible, typically undergoes hypertrophic differentiation late in embryonic development with central regions becoming resorbed by birth (Svandova et al., 2020). Meckel’s cartilage did not resorb in *C^KO^D^KO^* as compared to WT, indicating deficiencies in terminal chondrocyte differentiation (Fig. S5). To examine chondrocyte differentiation in more detail, we performed IF on sagittal sections through the cranial base. Examination of the PS region derived from cNCC origins demonstrated a loss of H3K4me1 in both *D^KO^* and *C^KO^D^KO^*in proliferative and hypertrophic chondrocytes, similar to observations in the tibia (Figure 3c-d, Figure 1b). SOX9 is appropriately activated in chondrocytes within *C^KO^D^KO^* synchondroses, but fails to be downregulated during hypertrophic differentiation (Fig. 3e), similar to observations in tibial growth plates. RUNX2 is activated in pre-hypertrophic chondrocytes; however, downstream targets such as type X collagen (COL10) exhibit minimal induction, supporting the previous observation that loss of KMT2C/D in chondrocytes prevents terminal hypertrophic differentiation (Fig. 3e).

### KMT2C/D deletion in stem cells restricts initial chondrocyte specification

As *Col2a1-Cre* deletes KMT2C/D after the chondrocyte has been established from progenitor stem cells, we sought to identify if these methylases may be functional in earlier steps of chondrocyte specification. To do so, we drove *Kmt2c/d* conditional knockout with a *Sox10-Cre* transgene (Matsuoka et al., 2005) active within cNCC stem cells of the anterior cranial base (Fig. 1a). Similar to *Col2-Cre* driven knockout, loss of both methylases in *C^KO^D^KO^-S10* (*Kmt2c^fl/fl^ Kmt2d^fl/fl^ Sox10-Cre*) pups at birth eliminates PS bone formation and reduces BS bone dimensions. Posterior mesoderm lineages such as BO were unaffected by *C^KO^D^KO^-S10* mutation. In a distinct fashion, KMT2D SKO in cNCCs (*D^KO^-S10*) had a more significant reduction in PS bone dimensions compared to results utilizing *Col2-Cre* indicating that an earlier loss of KMT2D in the progenitor state may have a greater impact on bone formation than subsequent KMT2D loss in chondrocytes (Fig. 3f-g).

To examine the chondrocyte cellular state following methylase deletion in cNCC stem cells, we performed IF for resting and differentiated chondrocyte markers following *Sox10-Cre* deletion. In *D^KO^-S10* newborn pups, SOX9 positive chondrocytes are specified with appropriate expression of type II collagen (COL2). However, there is also a loss of COL10A1 indicating lack of terminal hypertrophic differentiation despite the presence of KMT2C in these chondrocytes (Fig. 3h). By comparison, *C^KO^D^KO^-S10* mice lacking both methylases express minimal levels of SOX9 but do not express COL2, indicating a deficiency in proper chondrocyte specification. Moreover, RUNX2 and COL10 expression were absent with loss of both KMT2C and KMT2D in these osteochondral progenitors indicating a failure to initiate hypertrophic differentiation. KMT2C/D appear to not only serve a role in the transition to hypertrophic differentiation but are also critical during chondrocyte lineage specification.

### KMT2C/D are required for chondrocyte specification and hypertrophic differentiation *in vitro*

To better study the temporal dynamics of KMT2C and KMT2D function in gene activation and histone modification, we switched to an *in vitro* cellular model of cNCC chondrocyte differentiation (Fig. 1a). We utilized an established primary cNCC stem cell line isolated from E8.5 mouse embryos that retains the potential to differentiate into all cNCC lineages (Ishii et al., 2012) and features an osteochondral expression profile similar to endogenous cNCCs (Musa et al., 2024). We transfected the cNCCs with a GFP/Cas9/*Kmt2d*-gRNA construct targeting a region similar to the location of the mouse floxed allele followed by single cell cloning and sequencing to identify two independent lines carrying trans-heterozygous frameshift mutations of *Kmt2d* (SKO) (Fig. S1). To generate DKO lines for KMT2C/D, we transfected SKO lines with *Kmt2c* CRISPR constructs. SKO and DKO lines demonstrated similar severe reductions in H3K4me1 with a mild loss of H3K4me2 and no observable changes in H3K4me3 (Fig. 4a, Fig. S6a). Therefore, KMT2D is the predominant H3K4 mono-methylase in these cNCC lines despite similar levels in KMT2C expression. Mild H3K4me2 phenotypes and lack of any marked H3K4me3 effect may be due to compensation by other H3K4 methyltransferases such as KMT2A/B or KMT2F/G (SET1A/B). These findings validate that the knockout of KMT2D impacts the methylation status of H3K4.

**Figure 4:**
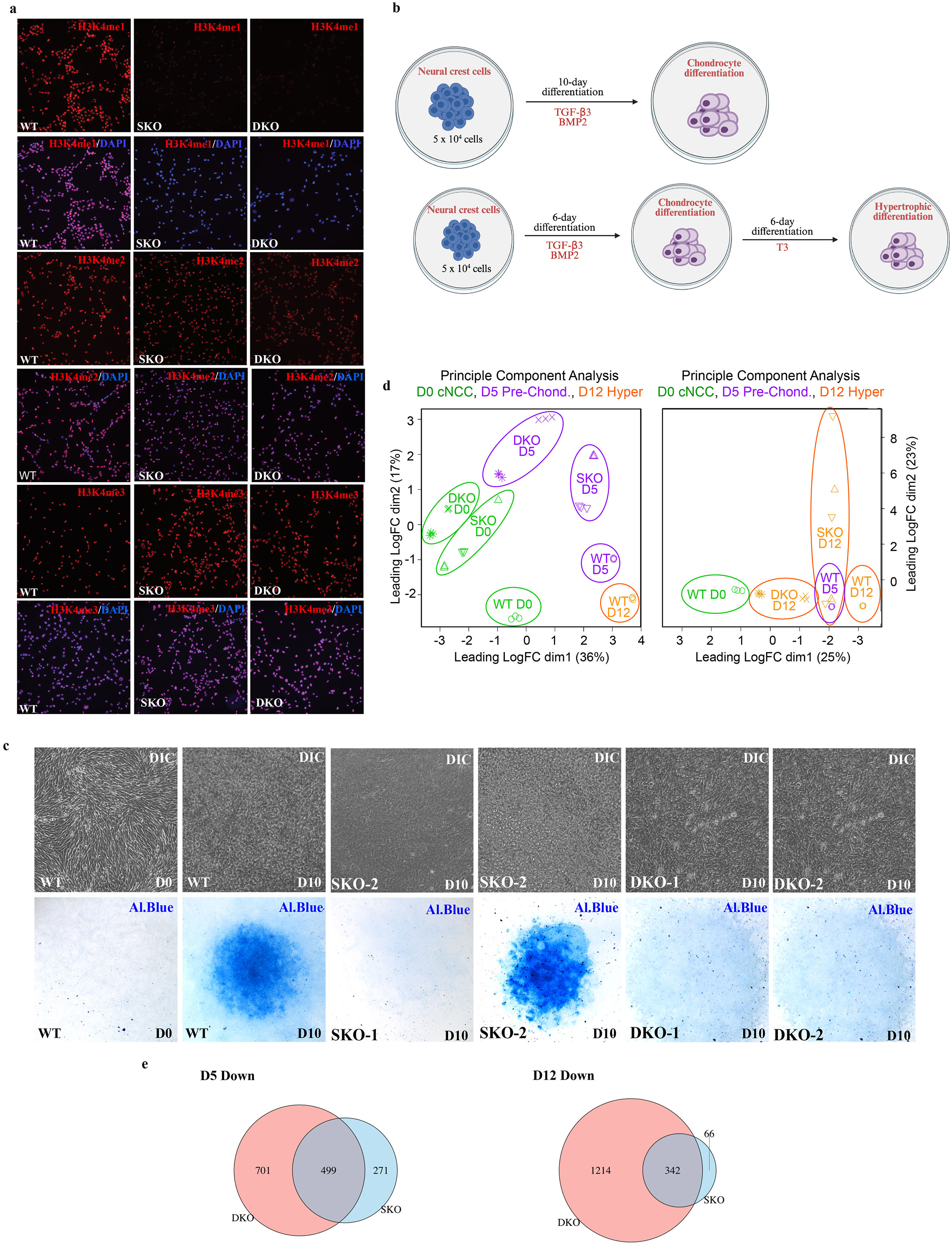
KMT2C/D are required for chondrocyte specification *in vitro*. **a** Immunofluorescence for H3K4me1, H3K4me2, and H3K4me3 in WT, SKO, and DKO lines with or without DAPI. **b** Differentiation model for NCC in vitro culture depicting 10-day chondrogenic differentiation and 12-day hypertrophic differentiation. **c** Alcian blue staining for proteoglycan production and DIC imaging of WT, SKO, and DKO lines at D0 (WT) and D10 of differentiation. **d** MDS plots of WT (circles), SKO line 1 (triangles), SKO line 2 (inverted triangles), DKO line 1 (x), and DKO line 2 (asterisks) in D0 of cNCCs (green), D5 of chondrogenesis (purple), and D12 of hypertrophic differentiation (orange). **e** Venn diagram of down-regulated genes in SKO and DKO lines as compared to WT at D5 and D12 of differentiation.

To differentiate cNCCs in culture, we used a previously established micromass plating method and incubated cells in culture for 10 days in chondrogenic media containing BMP2 and TGF-β3 (Fig. 4b) (Stanton et al., 2004; Underhill et al., 2014). At day 10, alcian blue staining of WT micromasses showed robust staining for chondrocyte proteoglycans, whereas SKO lines had variable staining with the majority of SKO lines (2/3) stained positively for alcian blue (Fig. 4c). In contrast, DKO lines consistently lacked alcian blue staining at 10 days of differentiation (2/2 lines). Together, these observations demonstrate that loss of KMT2C/D prevents chondrogenic cell type specification *in vitro*.

To emulate hypertrophic differentiation, we cultured cNCC micromasses for 6 days in chondrogenic media and an additional 6 days in hypertrophic media containing T3 (Mueller & Tuan, 2008). We performed bulk RNA-sequencing and differential expression analysis on WT, SKO, and DKO cNCC-derived cultures in triplicate at undifferentiated (day 0 = D0), chondrogenic (day 5 = D5), and hypertrophic (day 12 = D12) time points. RT-PCR indicated that WT chondrogenic micromasses exhibited marked increases in COL2A1 (87-fold), IHH (54-fold), and SOX9 (4.4-fold) expression as previously described (Fig. S6b) (Haseeb et al., 2021; Vortkamp et al., 1996).These time points were selected to capture dynamic gene expression changes associated with early lineage commitment and proliferative expansion. We utilized 2 independent clones of SKO and DKO mutations each including the one SKO line that negatively stained for alcian blue during differentiation. We found that undifferentiated WT cNCCs expressed both KMT2C and KMT2D and expression increased upon differentiation (Fig. S6c).

Multidimensional scaling (MDS) plots of RNA-seq data displayed progressive transcriptional divergence over the course of WT differentiation (Fig. 4d). By contrast, SKO D5 micromasses demonstrated an intermediate profile of differentiation. DKO D5 micromasses clustered most closely to undifferentiated (D0) WT samples, supporting the observation that loss of KMT2C/D impairs the initiation of the WT chondrogenic transcriptional program through redundant mechanisms. At D12 of hypertrophic differentiation, SKO lines have not advanced past the WT D5 chondrogenic profile (Fig 4d), supporting the *in vivo* data that KMT2D is required for hypertrophic differentiation. When comparing differential gene expression profiles of DKO to SKO, we observed some divergence in common downregulated mis-expressed genes at D5 of chondrogenic differentiation, supported by the MDS plot (Fig. 4e). Interestingly, there was a convergence of common SKO and DKO downregulated genes at D12 of hypertrophic differentiation indicating that neither genotype is capable of hypertrophic differentiation. These data are consistent with *in vivo* observations in *Col2a1-Cre* and *Sox10-Cre* mice that loss of KMT2D alone impacts terminal differentiation, whereas loss of both KMT2C/D affect both chondrocyte specification and hypertrophy.

### Loss of KMT2C/D disrupts RUNX2-and SOX9-driven gene programs governing skeletal system development

To assess molecular pathways leading to disrupted bone formation in KMT2C/D mutant mice, we performed Gene Set Enrichment Analysis (GSEA) on a ranked list of all expressed genes from our *in vitro* RNA-seq. This approach avoids the use of arbitrary thresholds, such as those used in edgeR, by ranking genes based on log_2_ fold-change (log_2_FC) without prefiltering. We analyzed enrichment against MSigDB C5 GO:BP gene sets containing terms related to skeletal development and endochondral ossification. Significance was determined using normalized enrichment scores (NES) and false discovery rate (FDR)-adjusted p-values (q-value). In DKO cells, GSEA revealed significant negative enrichment of the “Skeletal System Development” and “Ossification” gene sets at D5 and D12 of differentiation (NES < -1.6, q-value < 0.01) (Fig. 5a). The leading-edge genes (those driving the enrichment signal) were associated with key skeletal processes, including cartilage proliferation and terminal differentiation, indicating that KMT2C/D loss disrupts critical regulatory networks during bone formation.

**Figure 5:**
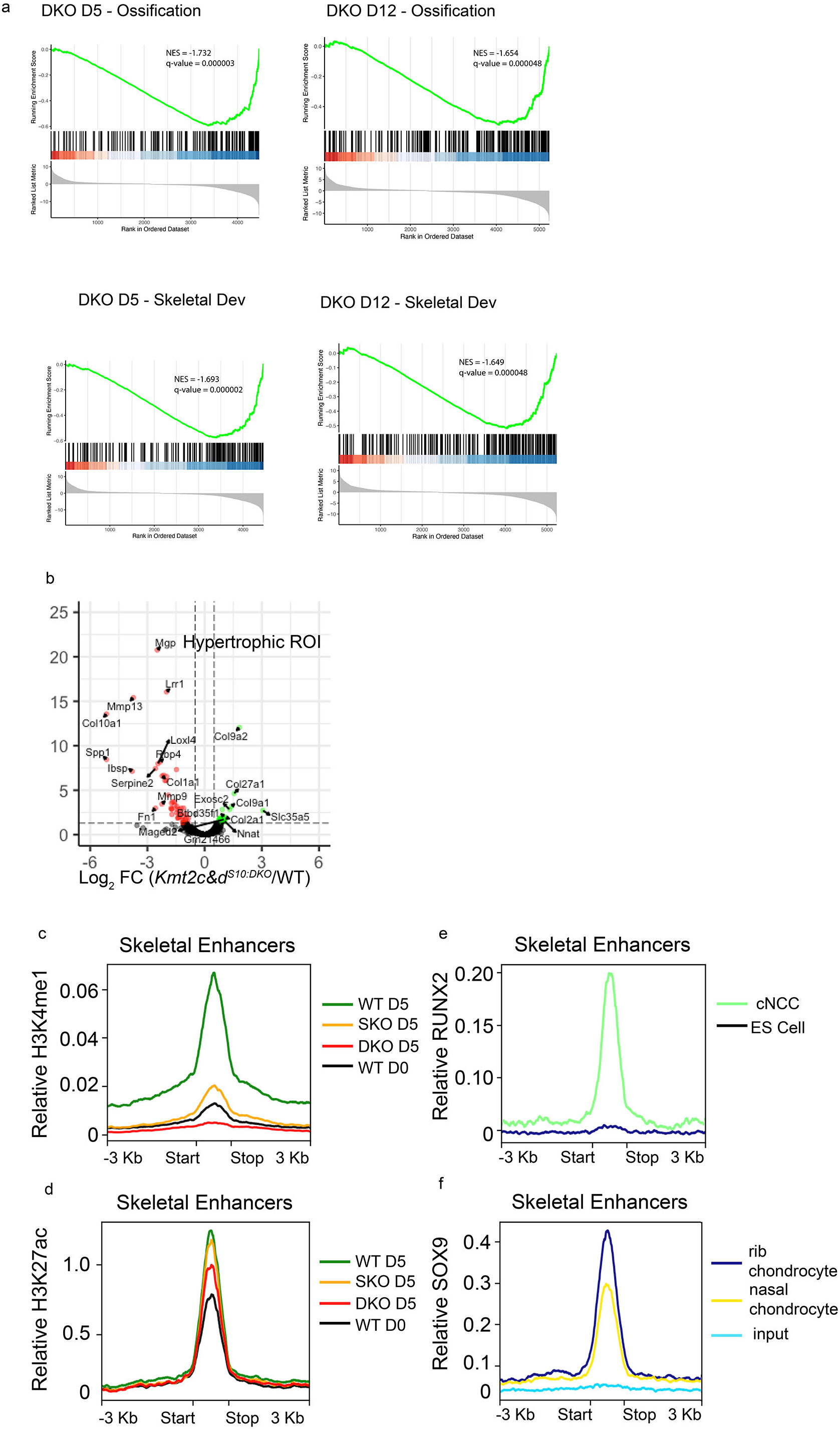
KMT2C/D loss impairs enhancer activity at key regulators of skeletal system development. **a** GSEA of all dysregulated genes in DKO as compared to WT at D5 or D12 of differentiation for “Ossification” and “Skeletal System Development” gene sets. **b** Volcano plot of spatial transcriptomics analyses illustrating log2FC of *C^KO^D^KO^-S10* differentially expressed genes from the hypertrophic regions relative to WT. Genes with log2FC < 1 or > -1 are demarcated in green or red, respectively. **c-d** Plot profile of H3K4me1 or H3K27ac levels at WT D5 enhancers associated with downregulated genes in DKO D5 linked to skeletal development (from part a). Illustrated are respective levels at WT D0 (black), WT D5 (green), SKO D5 (orange), and DKO D5 (red). ± 3 Kb up-and downstream of enhancer peaks depicted. **e-f** RUNX2 and SOX9 occupancy at enhancers associated with downregulated genes in DKO D5 linked to skeletal development (from part a). Illustrated in part e are RUNX2 CUT&RUN in D0 cNCCs (green) and ES cells (black). Illustrated in part f are SOX9 ChIP-seq from rib chondrocytes (dark blue), nasal chondrocytes (yellow), and input controls (light blue).

To identify molecular mechanisms whereby KMT2C/D promote chondrocyte specification, we focused on SOX9-regulated pathways and targets given the lack of COL2 expression in chondrocytes derived from mutant embryonic cNCCs (Fig. 3h). DKO lines showed significant downregulation of SOX9 targets including *Acan* and *Col2a1*, at both D5 and D12 of differentiation, along with a reduction in *Sox9* expression itself. Again, SKO lines exhibited more mild changes. These transcriptional changes suggested that KMT2C/D loss could disrupt upstream regulatory elements affecting SOX9 activity and expression, prompting us to investigate alterations in enhancer activity that could underlie the observed changes to gene expression.

Many KMT2C/D dysregulated genes in the GSEA analyses for skeletal development are direct RUNX2 targets. We selected a subset (∼10 genes) for further analysis across our *in vitro* RNA-seq datasets. RUNX2 targets such as *Ibsp*, *Col1a1*, *Col1a2, Mmp9*, and *Mmp13* were significantly downregulated in both DKO lines at D5 and D12 of differentiation. SKO lines showed milder gene dysregulation, correlating with less severe differentiation phenotypes. RUNX2 expression itself was also reduced, particularly at D12, suggesting compromised activation of its downstream gene program.

To validate the in vitro results indicating that disrupted RUNX2 regulation may be the cause of deficient skeletal development in KMT2C/D mutant mice, we performed spatial transcriptomic analysis in WT, *D^KO^-S10*, and *C^KO^D^KO^-S10* cranial bases at birth, focusing on the resting, proliferative, and hypertrophic zones of the PS region (Fig. S7a-b). These zones were delineated by a combination of H&E and nuclear stained cellular morphology as well as RUNX2 and COL10 IF. Although the assay was unable to detect many differences in resting and proliferative zones, *D^KO^-S10* and *C^KO^D^KO^-S10* hypertrophic chondrocytes showed dysregulated expression of multiple RUNX2-regulated genes such as *Col10a1*, *Mmp13*, *Mmp9*, and *Ibsp* (Fig. 5b). GSEA of these spatially defined transcripts also revealed significant negative enrichment of “Skeletal System Development” and “Ossification” gene sets, consistent with our *in vitro* findings (Fig. S7c).

### KMT2C/D loss impairs enhancer activity at key regulators of skeletal development

KMT2C and KMT2D are predominantly responsible for H3K4 mono-methylation in cNCCs and embryonic chondrocytes (Fig. 1b, Fig. 3c). To identify direct genomic targets of KMT2C/D, we performed CUT&Tag profiling of H3K4me1 in D0 cNCCs and D5 chondrocytes in WT, SKO, and DKO lines. H3K4me1 WT peaks of enrichment were dramatically lost at a large set of cNCC targets in both SKO and DKO cells (Fig. S8). The majority of these peaks were not located at transcription start sites, but rather at putative enhancers. To identify whether KMT2C/D regulated enhancers were active, we performed CUT&Tag for histone acetylation (H3K27ac). KMT2C/D regulated enhancers were active in WT cNCCs with H2K27ac enrichment. SKO and DKO cNCCs exhibited a similar reduction but not absence of enhancer activation. Notably, a smaller set of H3K4me1 peaks were gained specifically in DKO cNCCs at promoters.

To identify potential enhancer-gene regulatory interactions, we applied the Activity-By-Contact (ABC) model, integrating publicly available Hi-C data from wildtype mouse NCCs with our H3K27ac CUT&Tag datasets from D0 and D5 WT and DKO lines. We retained only strong enhancer-promoter pairs (ABC score > 0.02) for downstream analysis. Focusing on genes regulating “Ossification” and “Skeletal System Development” (GSEA Fig. 5a) that lose expression in DKO at D5 of chondrocyte differentiation, we identified 82 genes linked to 1,030 unique putative enhancers. These enhancers showed a significant gain in H3K4me1 and H3K27ac between D0 and D5 in WT, consistent with expected activation during chondrocyte differentiation (Fig. 5c-d). By contrast, DKO D5 samples failed to exhibit any H3K4me1 and showed markedly reduced H3K27ac at the same sites. SKO D5 samples displayed low levels of H3K4me1 and intermediate H3K27ac levels compared to DKO, albeit not to WT levels, suggesting partial retention of enhancer activation capacity due to compensation by KMT2C. We conclude that KMT2C and KMT2D are redundantly priming and activating enhancers to regulate chondrogenic gene expression critical for bone formation.

Given our prior observation that multiple downstream targets of RUNX2 and SOX9 were mis-regulated in SKO and DKO lines, we assessed direct TF occupancy at these enhancers. Using our prior RUNX2 CUT&RUN data from WT D0 NCCs and publicly available SOX9 ChIP-seq data from mouse rib and nasal chondrocytes (Ohba et al., 2015), we evaluated binding enrichment at our identified enhancers linked to KMT2C/D regulated skeletal development (Fig. 5e-f). Both RUNX2 and SOX9 showed significant enrichment at these sites relative to negative controls, consistent with direct regulatory roles. These results suggest that KMT2C/D loss may impair enhancer activation not only by reducing chromatin accessibility and H3K27ac deposition, but also by coordinating SOX9 and RUNX2 occupancy or regulation at key enhancers driving endochondral ossification.

## Discussion

Prior literature has established a partially redundant role for KMT2C and KMT2D in adipose tissue development, although this redundancy remains unexplored in skeletal development (Hu et al., 2013; Jang et al., 2019; J.-E. Lee et al., 2013). Accordingly, we generated chondrocyte-specific and cNCC-specific knockout mouse models for *Kmt2c* and *Kmt2d*, establishing partially redundant roles for these epigenetic regulators in the stage-specific progression of chondrocyte differentiation. Previous studies show that loss of KMT2D and UTX in MSCs impairs chondrocyte and osteoblast differentiation *in vitro*, suggesting a broader role for KMT2D beyond craniofacial development (Fasciani et al., 2020; Hemming et al., 2014). Expanding on these findings, we show that KMT2C and KMT2D are critical for endochondral ossification from MSC-and cNCC-derived chondrocytes. Consistent with prior observations, our results also indicate that KMT2C and KMT2D are not functionally equivalent in mammalian development (Kobayashi et al., 2024). Chondrocyte-specific deletion of both *Kmt2c* and *Kmt2d* or *Kmt2d* alone leads to marked skeletal defects, including delayed development and reduced bone formation, whereas deletion of *Kmt2c* alone produces no significant abnormalities in murine skeletal development or microarchitecture. Notably, combined loss of both methyltransferases results in more severe impairments to cartilage and bone formation than loss of either gene alone. These findings underscore a more prominent role for KMT2D in skeletal development while also revealing partial functional redundancy, as a single allele of *Kmt2c* in *D^KO^*partially rescues the *C^KO^D^KO^* phenotype and a single allele of *Kmt2d* in *C^KO^* preserves wild-type skeletal morphology. Collectively, we provide evidence that supports a broader role for KMT2C and KMT2D in skeletal development beyond a craniofacial context and suggests that disruption of epigenetic regulators may contribute to a wider spectrum of undiagnosed skeletal dysplasia.

In Shpargel et al., 2020, we showed that loss-of-function of KMT2D in cNCCs results in an inability of craniofacial chondrocytes to enter the hypertrophic stage of differentiation. Here, we expand on this observation in the chondrocyte lineage, showing that deletion of both KMT2C and KMT2D subsequent to chondrocyte specification impairs hypertrophic differentiation. This has profound effects on later skeletal formation and patterning, offering another potential mechanism behind delayed ossification or mineralization observed in pediatric growth disorders. We also find that loss of both KMT2C and KMT2D prevent earlier chondrocyte lineage specification, evidenced by lack of expression of COL2A1 in the *Sox10-Cre* model and an inability of DKO cNCCs to differentiate *in vitro*. As cNCC-derived tissues are impacted in craniofacial skeletal development disorders, these observations link early developmental gene regulation during chondrocyte lineage specification to downstream structural anomalies characteristic of KLEFS2 and KS.

Prior literature has shown that promoter-bound KMT2D upregulates *Shox2* within differentiated teratocarcinoma cells, allowing it to inhibit *Sox9* to restrict chondrogenesis (Fahrner et al., 2019). *Sox9* expression is upregulated in SKO and DKO cNCCs. However, we do not see dysregulation of *Shox2* in cNCC derived SKO or DKO lines in our bulk RNA-seq data prior to or during differentiation, suggesting that *Sox9* regulation in our *in vitro* model is *Shox2*-independent. We observe that *Sox9* expression does become reduced during in vitro differentiation in the absence of KMT2C/D. By contrast, *in vivo*, we observe continued expression of *Sox9* in *C^KO^D^KO^*pre-hypertrophic chondrocytes at E15.5, implying a failure to transition to RUNX2-dominant hypertrophy, consistent with impaired terminal differentiation. Previously, it has been shown that SOX9 directly interacts with RUNX2 to prevent its ability to access target genes (Zhou et al., 2006). Accordingly, excess SOX9 may prevent RUNX2-driven gene activation in the absence of KMT2C/D. It is unclear why there is a divergence between *in vivo* and *in vitro Sox9* expression patterns in the absence of KMT2C/D. Extrinsic cues *in vivo* may sustain *Sox9* expression whereas micromass cultures lack these contextual signals, subsequently affecting chondrocyte identity. Additionally, cNCCs lack KMT2C/D prior to chondrogenic lineage commitment *in vitro,* whereas in the *Col2a1-Cre* mouse, gene deletion occurs after specification (Fig. 1a). Despite this discrepancy, we observe similar downregulation of *Runx2* and downstream target genes such as *Col10a1* in *in vitro* differential expression analysis, *Sox10-Cre* spatial transcriptomic data, and *Col2a1-Cre/Sox10-Cre* immunofluorescence. These findings support the idea that absence of KMT2C/D disrupts the transition from SOX9 to RUNX2-guided chondrocyte differentiation, although further studies are necessary to define direct gene regulation or enhancer dysfunction as causative factors.

At the molecular level, prior studies have examined KMT2D function by assaying promoter H3K4me3; however, KMT2D lacks trimethylation activity *in vitro* (Zhang et al., 2015). Our *in vitro* observations support this finding as we do not see altered levels of H3K4me3 in SKO or DKO lines; however, we do observe global loss of H3K4me1 and a reduction in H3K4me2 levels upon loss of KMT2D or both KMT2C and KMT2D. In addition to a loss of H3K4me1 at enhancers, we also see a reduction in H3K27ac pointing to enhancer activation impairment. This epigenetic disruption corresponds with an inability of DKO lines to properly differentiate. Further investigation of downregulated skeletal genes in DKO lines revealed that associated putative enhancers fail to gain H3K4me1 and H3K27ac upon differentiation, which are normally acquired in WT chondrocytes. SKO lines retain partial enhancer activity, consistent with partial compensation by KMT2C, as shown by both milder chromatin and gene expression effects. Using WT RUNX2 CUT&RUN data and publicly available SOX9 ChIP-seq data, we found that RUNX2 and SOX9 binding was enriched at these ABC-predicted sites, supporting their role as functionally active regulatory elements in the skeletal gene expression program. These findings support a model in which KMT2C/D alter the active enhancer landscape during critical transition points of chondrocyte differentiation to allow for appropriate regulation of downstream skeletal gene regulation.

We provide comprehensive *in vivo* and *in vitro* models to determine the roles of KMT2C and KMT2D at two critical transition points, both chondrocyte specification and hypertrophy, in chondrocyte differentiation and endochondral ossification. The results of this study point to epigenetic enhancer dysfunction as a mechanism for skeletal malformations and provide new evidence for unequal partial redundancy between KMT2C and KMT2D in skeletal development. Differing outcomes from the *Col2a1-Cre* and *Sox10-Cre* deletions emphasize the importance of timing and chondrocyte lineage specificity in KMT2C/D function. Future studies will utilize primary mouse MSC cultures differentiated into chondrocytes to help clarify lineage-specific roles of KMT2C/D and allow direct assessment of SOX9 and RUNX2 chromatin binding dynamics, potentially resolving discrepancies observed between cNCC and MSC-derived models of KMT2C/D deficiency in endochondral ossification. Together, we provide evidence for regulation of chondrocyte differentiation by KMT2C/D modulation of enhancer activity, highlighting the need for future research in epigenetic regulators in skeletal development. Such studies may continue to uncover how disruptions in enhancer activity contribute to skeletal dysplasia.

## Materials and Methods

### Mice

Mouse colony maintenance and experimentation conducted according to University of North Carolina at Chapel Hill Institutional Animal Care and Use Committee protocols. Mice crossed for at least 4 generations on C57BL/6J background prior to experimentation. *Col2a1-Cre* mice obtained from Dr. Richard Loeser (Ulici et al., 2019) and originally developed by Dr. Di Chen (Rush University Medical Center, Chicago, IL) (Chen et al., 2007). *Sox10-Cre* mice were obtained from The Jackson Laboratory (Matsuoka et al., 2005). *Kmt2c^fl^* allele was previously described (Jang et al., 2019). *Kmt2d^fl^* allele was previously described (Jang et al., 2019; Shpargel et al., 2014, 2020). Cre-activated *Rosa^Tomato^* mice were obtained from The Jackson Laboratory (Madisen et al., 2010). To avoid genetic background biases, all analyses were performed with Cre-negative littermates. For lineage analysis, *Rosa^Tomato^* mice were crossed to *Kmt2c^fl/fl^ Kmt2d^fl/fl^* females. *Kmt2c^fl/+^ Kmt2d^fl/+^ Col2a1-Cre* males and *Kmt2c^fl/fl^ Kmt2d^fl/fl^* females were crossed and the resulting progeny were used for analysis. As the *Sox10-Cre* transgene is active in the male germline, the *Sox10-Cre* allele was crossed into *Kmt2c Kmt2d* mice as described (Shpargel et al., 2020). Genomic DNA was prepared from tail clips and PCR analysis was used for genotyping of *Kmt2c^fl^*, *Kmt2d^fl^*, *Col2a1-Cre*, *Sox10-Cre,* and *Rosa^Tomato^* mice. All primers are available upon request.

### Tissue Preparation

For E15.5 collection, limbs were removed in 1x PBS on ice using a Leica MZFLIII microscope at 0.8X. Samples incubated in 4% PBS-buffered paraformaldehyde for 50 minutes at 4°C, washed 3 times in 1x PBS for 10 minutes, and processed in a sequential sucrose gradient at 4°C (10% sucrose/1x PBS, 15% sucrose/1X PBS, 20% sucrose/1X PBS). Samples incubated in 20% sucrose solution/Tissue-Tek O.C.T. Compound at a ratio of 1:1 for 2 hours at 4°C. Limbs were embedded in 20% sucrose/O.C.T. solution at a ratio of 1:3. Sectioning of 8-micron samples performed on Leica CM3050 S cryostat. For E18.5 and P0 mouse collection, mice were processed similarly to E15.5, but with an 80-minute initial fixation in 4% PBS-buffered paraformaldehyde at 4°C. These samples were collected at 10 microns. For collection of hindlimbs from 8-week-old mice, hindlimbs removed by hip dislocation method (Amend et al., 2016) keeping the knee joint intact. Additional tissue removed from limbs with scalpel and forceps. Limbs rinsed with 1X PBS and incubated at room temperature in 4% PBS-buffered paraformaldehyde for 48 hours. Limbs incubated for 6 days in 1X Immunocal (StatLab, 1414-1 GAL) to decalcify, refreshing solution at day 3, then washed 3 times in 1X PBS. Subsequent dehydration of limbs in gradient ethanol solutions. Samples processed by University of North Carolina at Chapel Hill Pathology Services Core using Citrisolv and paraffin processing method and embedded. Limbs repositioned in paraffin to section through knee joint coronally. Limbs sectioned in 10-micron sections on Leica RM2165 microtome.

### Histology

Eight-week limb samples deparaffinized using Citrisolv (Decon Labs, Inc., 1601), soaking 2 times for 5 minutes each. Samples were mounted using Citramount (Polysciences, 22253-120). Safranin O/fast green FCF staining performed as described (Bryan, 1955) using 0.02% fast green (3 minutes, Sigma-Aldrich, F7252), 1% acetic acid, and 0.2% safranin O (5 minutes, Sigma-Aldrich, 84120-25G) solutions. Toluidine blue O stain (5 minutes, Fisher Scientific, T161-25) performed as described(Schmitz et al., 2010). Hematoxylin (10 minutes, Harris Hematoxylin, Leica NC0338284) and eosin Y (1 minute, Fisher Scientific, E511-25) staining performed as described(Cardiff et al., 2014) with the following adjustments: After staining with Hematoxylin for 10 minutes, samples were dipped twice in dH_2_O to rinse, dipped twice in 0.02% acetic acid, dipped 10 times in dH_2_O to rinse, dipped 15 times in 0.015% ammonium hydroxide, dipped 10 times in dH_2_O, and stained in Eosin for 1 minute. For P0 skeletal staining, heads and limbs with skin removed were fixed in 95% ethanol and stained for Alizarin red S (Sigma-Aldrich, A5533) and alcian blue (Amresco, 33864-99-2) as described (Lufkin et al., 1992). Imaging conducted on Leica SP8X and Evident Slideview VS200 slide scanner. Measurements and image analysis conducted using FIJI/ImageJ (Schneider et al., 2012) or QuPath (Bankhead et al., 2017), respectively. Growth plate depth calculated across 160 µm, with 5 measurements taken every 40 µm for a total of 4 sections per limb. Proliferative and hypertrophic cell counts and areas were also calculated over 160 µm every 40 µm for a total of 4 sections per limb. Quantitation of longitudinal columns of proximal tibial growth plate chondrocytes was calculated using the FIJI/ImageJ multi-point tool to measure x and y spatial coordinates of each cell from the vertical origin point. Five columns down to a depth of ten cells beginning at the first identifiable proliferative chondrocyte were measured across 3 animals per genotype.

### Immunofluorescence

Immunofluorescence was performed in 1X PBS/3% BSA/10% goat serum/0.1% Triton-X-100 using SOX9, RUNX2 (1:150, Santa Cruz, sc-390351 or 1:800, Cell Signaling, 12556S), COLX (1:20, DSHB, X-AC9), OSX (1:150, Santa Cruz, sc-393325), BrdU (1:250, Abcam, ab6326 or 1:200, Cell Signaling, 5292S), UTX (1:200, Cell Signaling, 33510S), and H3K4me1 (1:400, Cell Signaling, 5326S). BrdU labeling was performed by maternal intraperitoneal injection at 50 mg/kg in 1X PBS 30 minutes prior to embryonic dissection (Sigma-Aldrich, B5002-250MG)(Musa et al., 2024). BrdU-labeled slides were treated with 2N HCl in 1X PBS at 37°C for 30 minutes before blocking. For hindlimb sections from 8-week-old mice, samples were deparaffinized and rehydrated then washed in dH_2_O twice. For antigen retrieval, samples were immersed in 1X ImmunoDNA Retriever with Citrate (Bio SB, BSB 0020) overnight at 55°C. Due to autofluorescent heme in 8-week samples, samples were treated with Vector TrueVIEW Autofluorescence Quenching Kit (Vector Laboratories, SP-8400). Cells in culture were fixed in 4% PBS-buffered paraformaldehyde at room temperature for 10 minutes, extracted for 5 minutes in 0.5% Triton-X-100/1X PBS, and blocked in 10% goat serum in 1X PBS before antibody incubation (30 minutes at 37°C).(Musa et al., 2024) Imaging performed with Leica SP8X. Images analyzed with FIJI/ImageJ(Schneider et al., 2012).

### Micro-computed tomography (microCT)

Following euthanasia, hindlimbs were removed and fixed in 4% PBS-buffered paraformaldehyde for 48 hours at room temperature. Micro-computed tomography analysis was performed using the Scanco Model 40 (Scanco Medical, Bassersdorf, Switzerland) scanner provided by UNC’s Biomedical Research Imaging Center. A voxel size of 12 microns was used and samples were scanned with a voltage of 70 kVp and a current of 114 uA. The integration time for each projection was 200 ms with a total of 500 projections per slab. To investigate KMT2C/D-associated changes in long bones, the trabecular and cortical architecture of the tibia were evaluated. Trabecular microstructure evaluation was performed in a 1.2mm region of interest beginning 0.12 mm below the tibial growth plate. Cortical measurements were assessed at the midshaft in a 1.2 mm region of interest beginning 2 mm below the growth plate. Global lower threshold values of 375 HA/cm^3^ for trabecular bone and 513.2 HA/cm^3^ for cortical bone were used. The relative bone volume (% BV/TV), trabecular number (Tb.N in 1/mm), trabecular thickness (Tb.Th in mm), trabecular separation (Tb.Sp in mm), and cortical thickness (Ct.Th in mm) were determined. To calculate relative bone volume, a voxel-based analysis was performed to calculate relative bone volume and a triangle-based surface reconstruction method was applied to analyze trabecular architecture. Representative three-dimensional micro-CT reconstructions generated using 3D Slicer (https://www.slicer.org) (Fedorov et al., 2012).

### Cell culture

O9-1 cranial neural crest cells were grown on Matrigel-coated (Corning, 356234, or Biotechne, 3432-005-01) plates in 50% mouse embryonic fibroblast (MEF)-conditioned DMEM and 50% growth media of the following: DMEM, fetal bovine serum (FBS), Penicillin/Streptomycin, L-glutamate, nonessential amino acids, sodium pyruvate, and β-mercaptoethanol supplemented with basic fibroblast growth factor 2 (FGF2), and leukemia inhibitory factor (LIF) to maintain NCC multipotency.(Ishii et al., 2012) To generate mutations, wildtype O9-1 cells were transfected with LentiCRISPRv2GFP (Addgene plasmid # 82416) containing gRNA targeting exon 49 in *Kmt2d*. We used the Crispor website (http://crispor.tefor.net) to identify the best guide recognition sequence. Clonal lines were screened to confirm frameshift mutations in both alleles to establish SKO-1, SKO-2, and SKO-3. We obtained trans-heterozygous mutations in exon 49 upstream of the loxP site found in *Kmt2d^fl^* mice. The same process was repeated in SKO-1 and SKO-2 lines using a gRNA targeting exon 56 in *Kmt2c* to establish DKO-1 and DKO-2. Similarly, we obtained trans-heterozygous mutations in exon 56 just upstream of the loxP site in *Kmt2c^fl^* mice.

### Chondrocyte differentiation

NCC cell lines were seeded using the micromass method(Stanton et al., 2004; Underhill et al., 2014) at a density of 5 x 10^4^ cells in a 10μl droplet and allowed to adhere for 2 hours before adding chondrogenic media (MEM α, Pen/Strep, sodium pyruvate, FBS, ITS, ascorbic acid, dexamethasone, TGF-B3, BMP2) as described(Ishii et al., 2012). Cells were cultured for 10 days in chondrogenic media, refreshing media every two days, before staining with alcian blue as previously described(Iezaki et al., 2019). To differentiate NCCs into hypertrophic chondrocytes, NCCs were seeded and cultured as described above in chondrogenic media for 6 days, then switched to hypertrophic media (MEM α, Pen/Strep, ITS, ascorbic acid, dexamethasone, triiodothyronine [T3], β-glycerophosphate) for an additional 6 days(Mueller & Tuan, 2008).

### Micromass dissociation

For micromass dissociation, cells were washed in 1x PBS then incubated in 2 mg/mL of type II collagenase (Worthington, LS004176). Micromass pellet was lifted off of the well using a cell scraper and solution containing the micromass was pipetted into a 2 mL tube. Samples were nutated at 37°C for 1 hour with a short vortex every 20 minutes. To neutralize collagenase, 10% FBS was added, and samples were spun down to pellet cells. Cells were transferred to a buffer of interest for any downstream applications (RNA-seq, CUT&Tag) to count cells.

### RNA-seq

RNA was isolated with Trizol (Thermo-Fisher) in three biological replicates from WT, DKO, or SKO undifferentiated NCC lines or cells at days 5 and 10 of chondrogenic or day 12 of hypertrophic differentiation. cDNA was synthesized, Truseq adapters were ligated, and library amplified with the KAPA mRNA HyperPrep Kit (Roche, 08098123702). RNA quality was assessed using Qubit fluorometer and Agilent Bioanalyzer. Library samples were multiplexed and sequenced on the NovaSeq 6000 platform using 50 bp paired-end sequencing. Analysis performed as described previously(Musa et al., 2024; Shpargel et al., 2020). Briefly, FastQC (http://www.bioinformatics.babraham.ac.uk/projects/fastqc/) was used to assess sequence read quality. Reads were aligned to the mm10 genome using Tophat2(D. Kim et al., 2013) RNA-seq reads were counted using htseq-count(Anders et al., 2015). EdgeR identified significant (FDR < 0.05) differential expression using DESeq2’s independent filtering method to determine minimum read count cutoffs per sample.(Anders & Huber, 2010; Robinson et al., 2010).

### CUT&Tag

To isolate cells for CUT&Tag (10^5^ cells), at least two biological replicates were collected from WT, DKO, or SKO NCC lines at the following stages: undifferentiated, days 5 and 10 of chondrogenic differentiation, or day 12 of hypertrophic differentiation. Cells were bound to Concanavalin A beads (Polysciences, 86057-3) and processed according to Epicypher pAG-Tn5 protocols (15-1017). Epicypher SNAP-CUTANA K-MetSat Panel (SKU 19-1002), a spike-in control specifically for histone post-translational modifications, was spiked into WT, DKO, and SKO H3K4me1 ConA-immobilized samples prior to primary antibody addition at a final dilution of 1:25 based on the initial starting number of cells. All antibody incubations and washing steps were carried out in PCR strip tubes, each containing 200 µl of solution with antibody dilutions as follows: IgG isotype control (1:100, Cell Signaling, 3900S), H3K27ac (1:100, Cell Signaling, 8173S) or H3K4me1 (1:100, Cell Signaling, 5326S). Tagmentation reaction products were treated with Proteinase K, followed by extraction, precipitation, washing and resuspension in 10 mM Tris/1 mM EDTA pH 8 (TE) buffer as described by Musa et al. 2024 (Musa et al., 2024) and the Epicypher CUT&Tag protocol. *Drosophila* SL-2 cells underwent a similar CUT&Tag protocol (cells were growth at room temperature in Schneider’s medium with 10% FBS) using H3K27me3 antibody (1:100, Cell Signaling, 9733S). These *Drosophila* H3K27me3 products were then spiked into WT, DKO, and SKO H3K27ac samples at a 1:10 ratio. NEBNext HiFi polymerase (New England Biolabs) was used to amplify libraries with dual-index primers that annealed to the Tn5 adapters as described (Buenrostro et al., 2015). Library samples were multiplexed and sequenced on the NovaSeq 6000 platform using 50 bp paired-end sequencing. FastQC (http://www.bioinformatics.babraham.ac.uk/projects/fastqc/) was used to assess sequence read quality. Reads were aligned to the mm10 or dm6 genome using Bowtie2 (Langmead & Salzberg, 2012) with the following options: -local –very-sensitive-local –no-unal –no-mixed –no-discordant – phred33 -I 10 -X 1500. Bam files were created with samtools view with the following options: -S -b -F 4 -q 30 (Li et al., 2009). PCR duplicated reads were removed with picard MarkDuplicates (http://broadinstitute.github.io/picard/). Significant enrichment was called with MACS version 2 after pooling replicates by using –broad -g mm –broad-cutoff 0.1. DeepTools computeMatrix, plotHeatmap, and plotProfile functions were used to generate heatmaps and profiles of H3K4me1 and H3K27ac read coverage at H3K4me1 peaks.

### Activity-By-Contact (ABC) model

Activity-By-Contact model as described (Bond et al., 2024). An adapted version of the ABC pipeline (Fulco et al., 2019) was used to identify enhancer-gene pairs using publicly available Hi-C data (GEO accession GSM6505198) derived from mouse cranial neural crest cells in wildtype E10.5 mandibular tissue (Kessler et al., 2023). Peaks were called from CUT&Tag H3K27ac bam files using MACS2 which were then used as the candidate enhancers for the ABC model. After peak calling, peaks were resized to 500 bp in size from the summit, mm10 blacklisted regions were removed, and overlapping peaks were merged. H3K27ac signal was extracted from candidate enhancers using bedtools (Quinlan & Hall, 2010). ABC scores were calculated between all genes and candidate enhancers within a 5 Mb window using mariner (version 1.2.1) (Davis et al., 2024). Enhancer-promoter pairs were called as valid if they had an ABC score > 0.02.

### Statistics

Experimental data were statistically analyzed using GraphPad Prism version 10.4.1 for Mac (Graphpad Software, Boston, Massachusetts USA, www.graphpad.com) using one-way ANOVA followed by Dunnett’s multiple comparisons test, Welch ANOVA followed by Dunnett’s T3 multiple comparisons test, or unpaired t-test with Welch’s correction. All data were expressed as mean ± standard deviation and a two-tailed Student’s t-Test was used for intergroup comparison. A score of P > 0.05 was considered statistically significant. Asterisks denote significant differences compared to wildtype (* = p-value < 0.05, ** = p-value < 0.01, *** = p-value < 0.001).

## Data Availability

RNA-seq and CUT&Tag datasets are in submission to FaceBase: https://doi.org/10.25550/8C-WDXG

## Acknowledgements

This work was financially supported by an NIH NIDCR R01 award (R01DE030530) and a NIH NIDCR R01 F31 award (F31DE033916).

## Supplemental

**S1:**
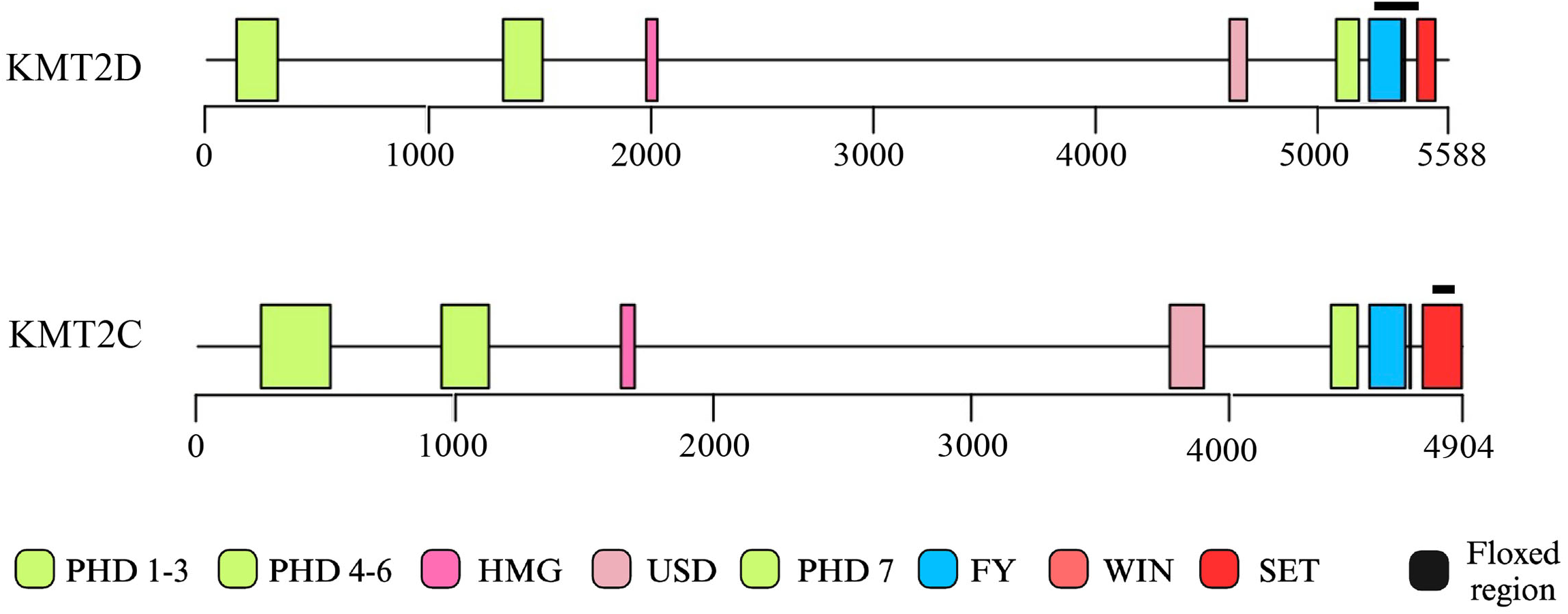
Schematic of KMT2C and KMT2D proteins with locations of PHD fingers (PHD), High Mobility Group domain (HMG), UTX Stabilization Domain (USD), FY-rich domain (FY), WW domain-interacting domain (WIN), and Suppressor of variegation, Enhancer of zeste, and Trithorax domain (SET). *In vivo* floxed regions annotated by the black bars (exon 50-51 in KMT2D and exon 56-58 in KMT2C). **S2:** Discoloration of *C^KO^D^KO^*pups as compared to WT at P0. **S3:** Micro-CT analysis of trabecular bone formation (Tb.Th, Tb.N) in 8-week-old tibia of WT, *C^KO^*, *D^KO^*, and *C^KO^D^KO^* mice (*N*=2, *N*=6, respectively). Data shown are mean ± SD. Statistical analyses were performed with ordinary one-way ANOVA followed by Dunnett’s test. *P<0.05, **P<0.01, ***P<0.001, ****P<0.0001, ns = not significant where specified

**S2:**
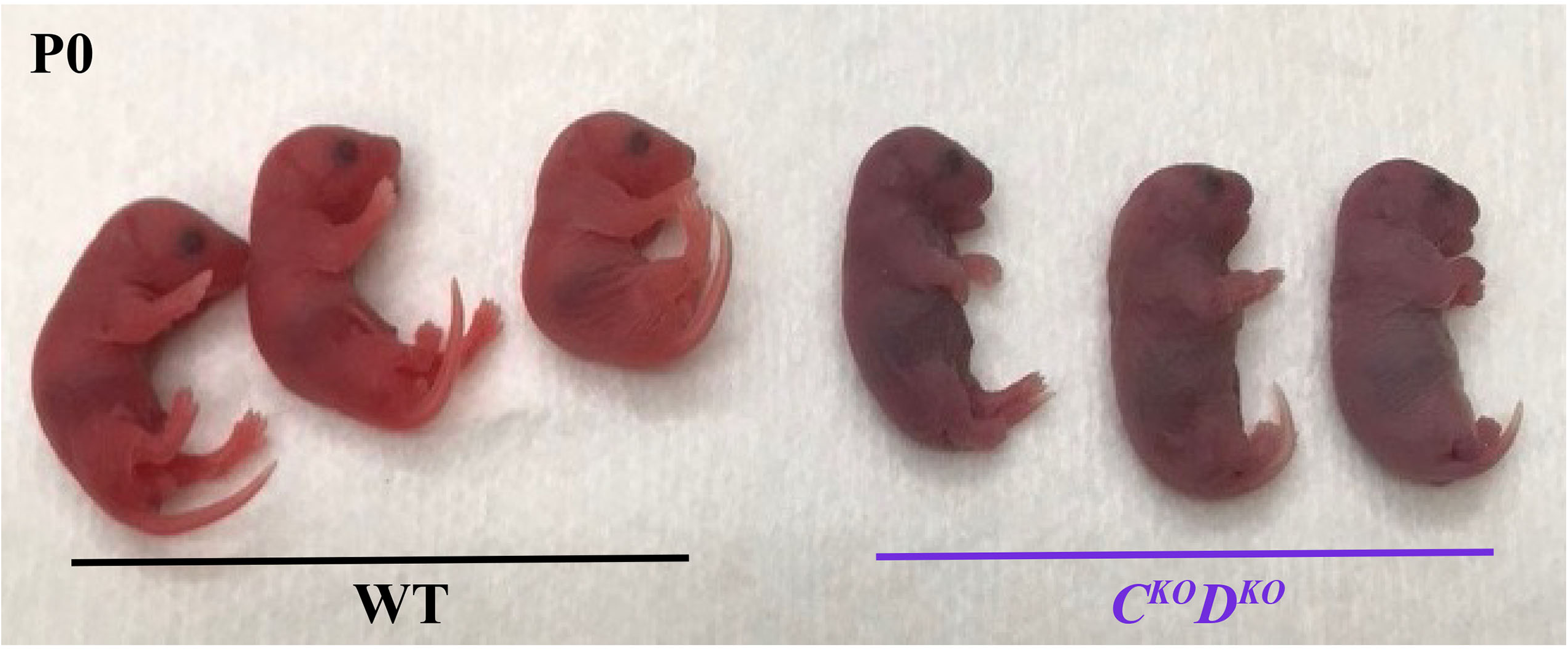
Discoloration of *C^KO^D^KO^* pups as compared to WT at P0.

**S3:**
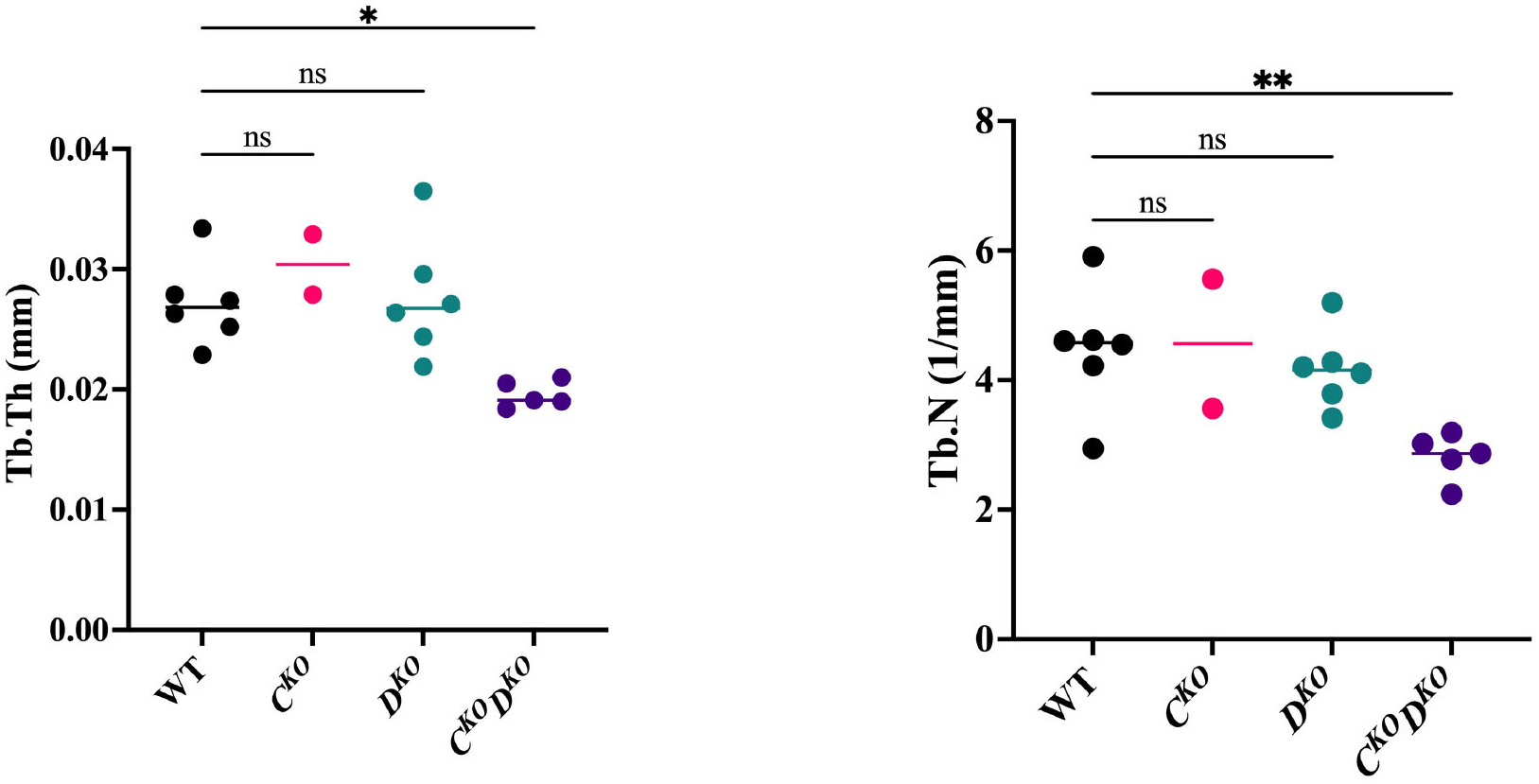
Micro-CT analysis of trabecular bone formation (Tb.Th, Tb.N) in 8-week-old tibia of WT, *C^KO^*, *D^KO^*, and *C^KO^D^KO^* mice (*N*=2, *N*=6, respectively). Data shown are mean ± SD. Statistical analyses were performed with ordinary one-way ANOVA followed by Dunnett’s test. *P<0.05, **P<0.01, ***P<0.001, ****P<0.0001, ns = not significant where specified

**S4:**
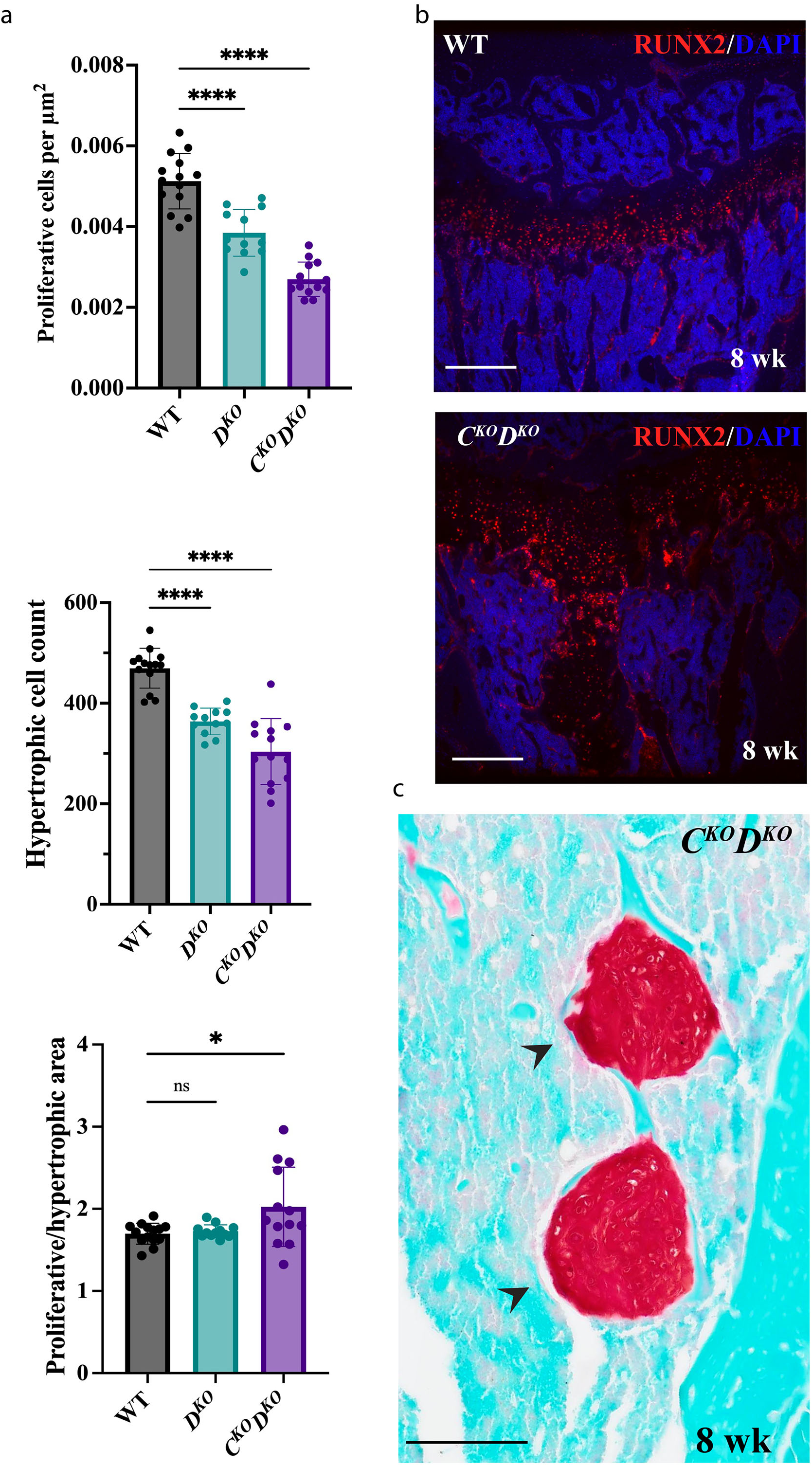
**a** The ratio of proliferative cells to proliferative area, proliferative to hypertrophic area, and the total number of hypertrophic cells per section in 8-week-old proximal tibial growth plates of WT, *D^KO^*, and *C^KO^D^KO^* (*N*=4, *N*=3, *N*=4, respectively). **b** Immunofluorescence of RUNX2 with DAPI in 8-week-old coronally sectioned proximal tibial growth plates of WT and *C^KO^D^KO^*indicating disorganization of *C^KO^D^KO^* growth plate and penetrance of RUNX2-expressing cells into the diaphysis. Scale bar is 250 µm. **c** Safranin O/fast green staining of the same region of RUNX2-expressing cells as shown in S4b *C^KO^D^KO^* section showing proteoglycan expression. Scale bar is 200 µm. Data shown are mean ± SD. Statistical analyses were performed using ordinary one-way ANOVA followed by Dunnett’s test *P<0.05, **P<0.01, ***P<0.001, ****P<0.0001, ns = not significant where specified

**S5:**
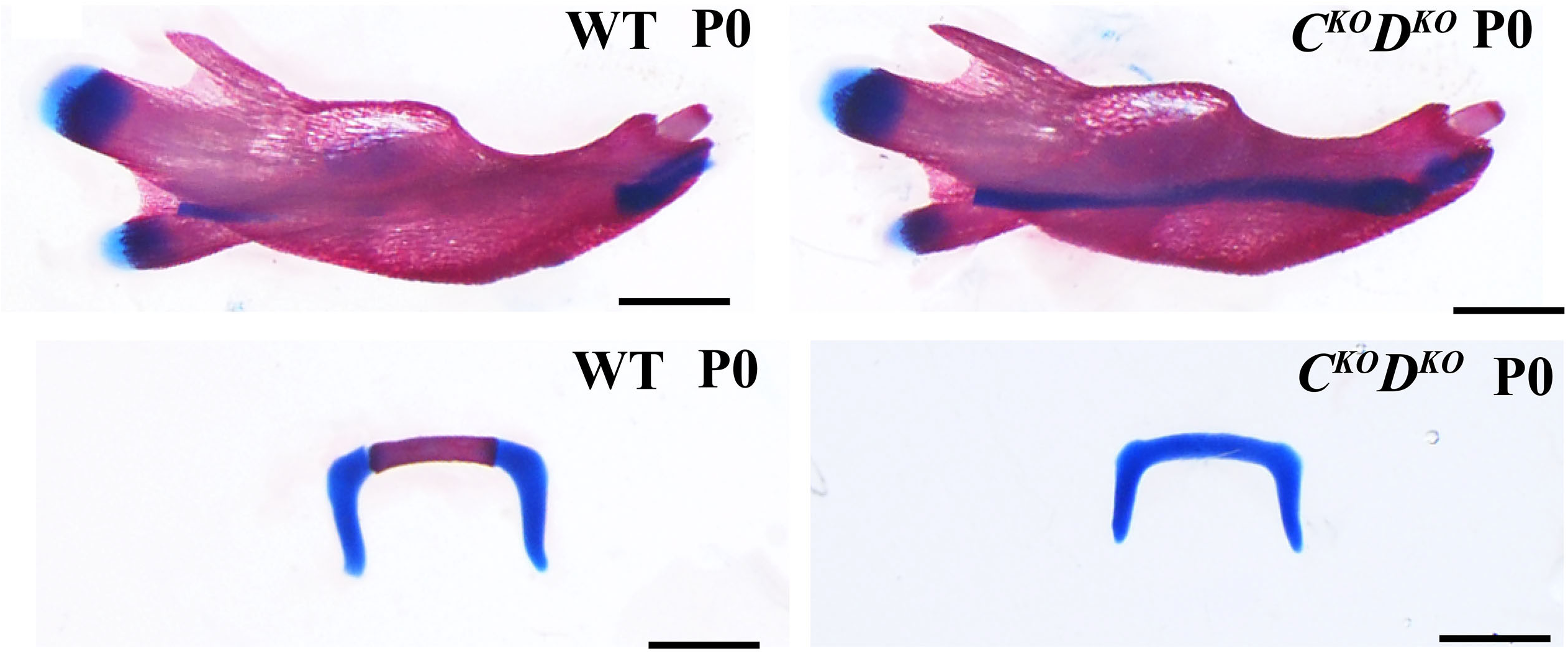
Wholemount images of alizarin red and alcian blue stained dissected mandible and hyoid cartilage in WT and *C^KO^D^KO^*P0 mice. Scale bar is 1 mm.

**S6:**
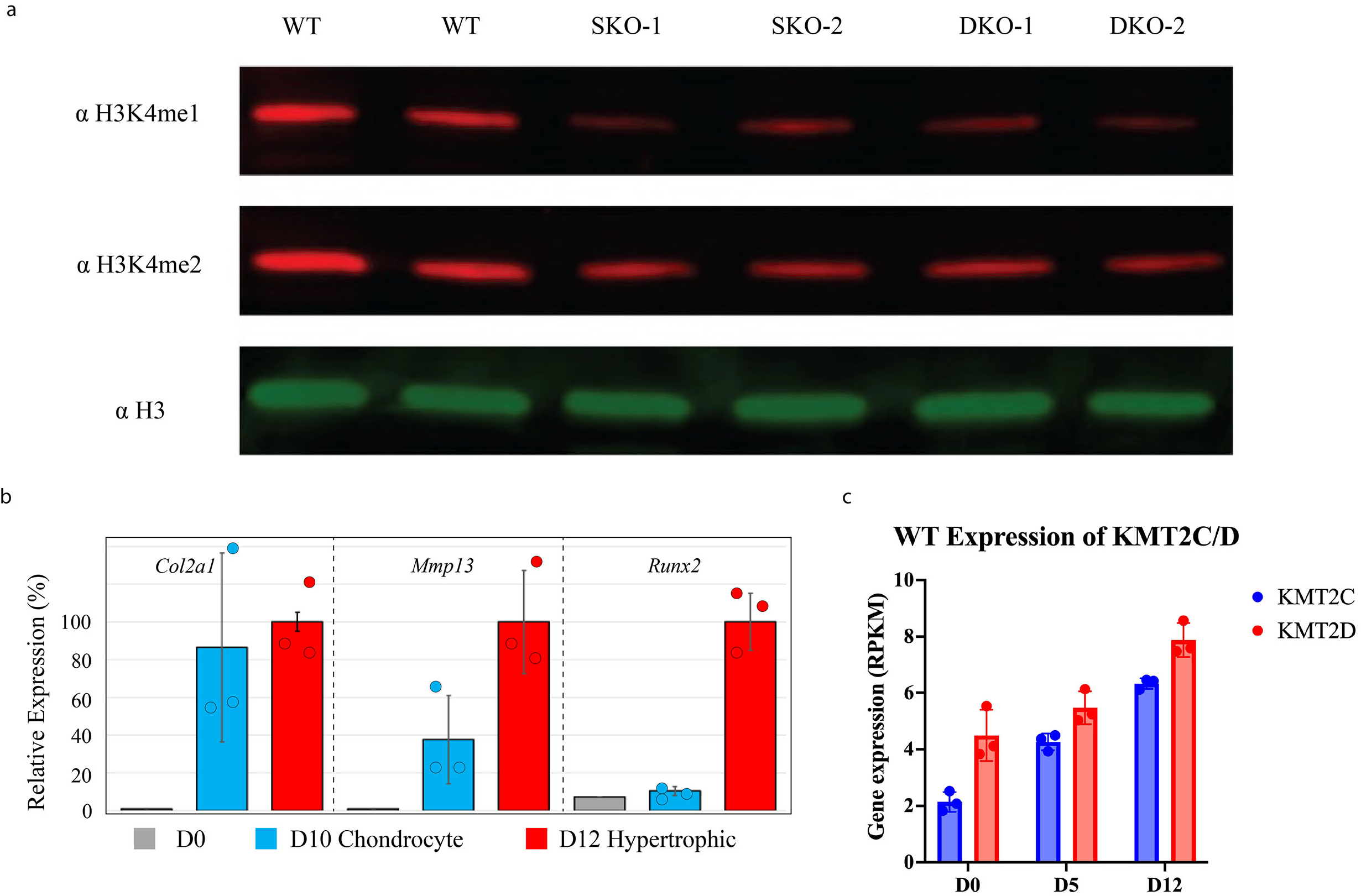
**a** Western blot for H3K4me1, H3K4me2, and H3 control antibodies in undifferentiated NCCs in WT, SKO line 1 (SKO-1), SKO line 2 (SKO-2), DKO line 1 (DKO-1) and DKO line 2 (DKO-2). **b** RT-PCR results of WT micromasses during chondrogenic differentiation at D0, D10, and D12 of differentiation. Showing relative expression as a percentage. **c** WT expression level (RPKM) of KMT2C and KMT2D at D0, D5, and D12 of chondrogenic differentiation. Data shown are mean ± SD.

**S7:**
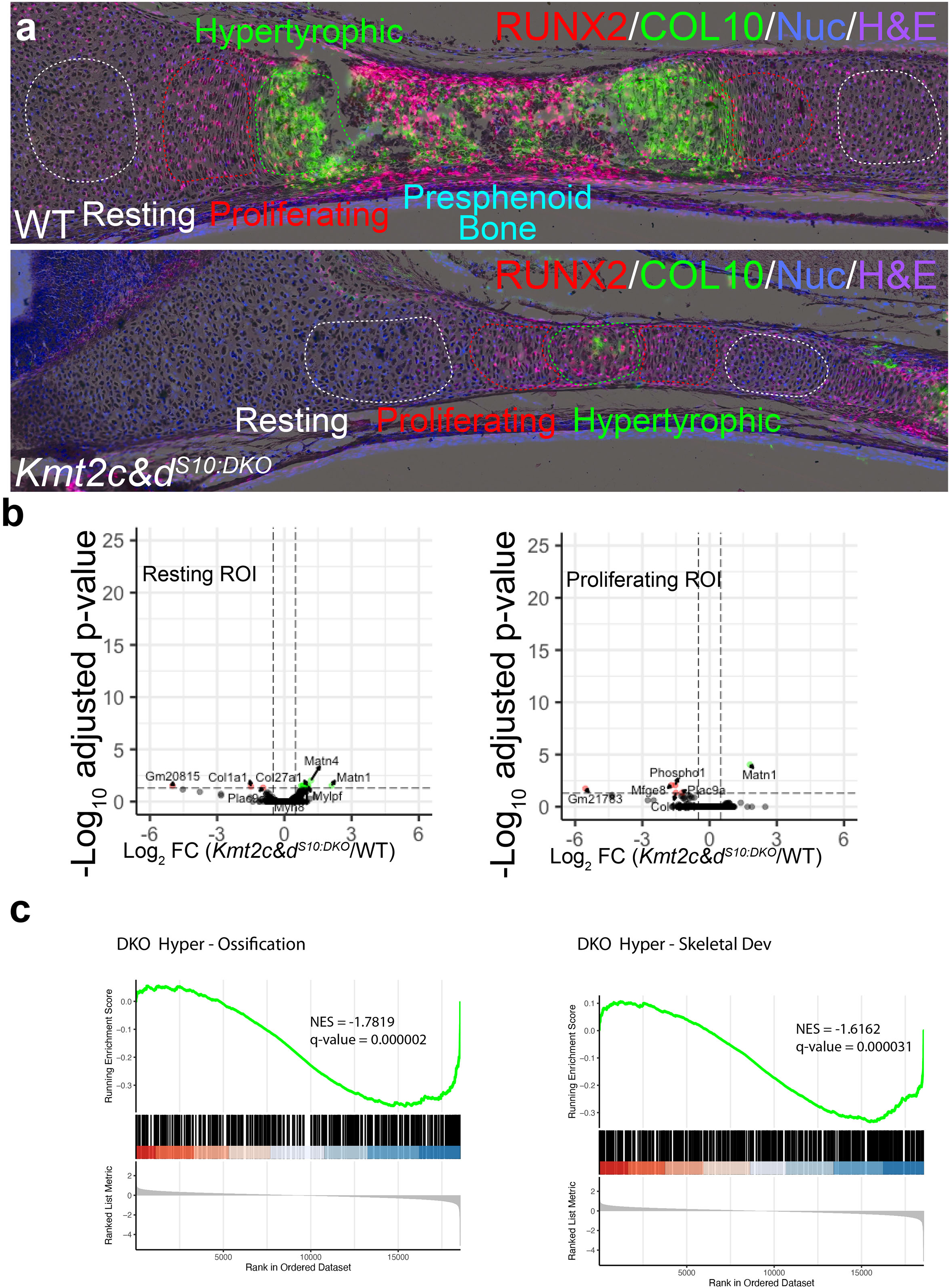
**a** Regions of sagitally sectioned presphenoid bone collected for spatial transcriptomic analysis. Hypertrophic (green), proliferative (red), and resting (white) regions collected for analysis demarcated by dashed circles. RUNX2 (red) and COL10 (green), nuclear staining (blue), and H&E staining shown. **b** Volcano plots of log2FC of *C^KO^D^KO^-S10* differentially expressed genes from the resting or proliferative regions relative to WT. Genes with log2FC < 1 or > -1 are demarcated in green or red, respectively. **c** GSEA of dysregulated hypertrophic region genes in *C^KO^D^KO^-S10* as compared to WT for “Ossification” and “Skeletal System Development” gene sets.

**S8:**
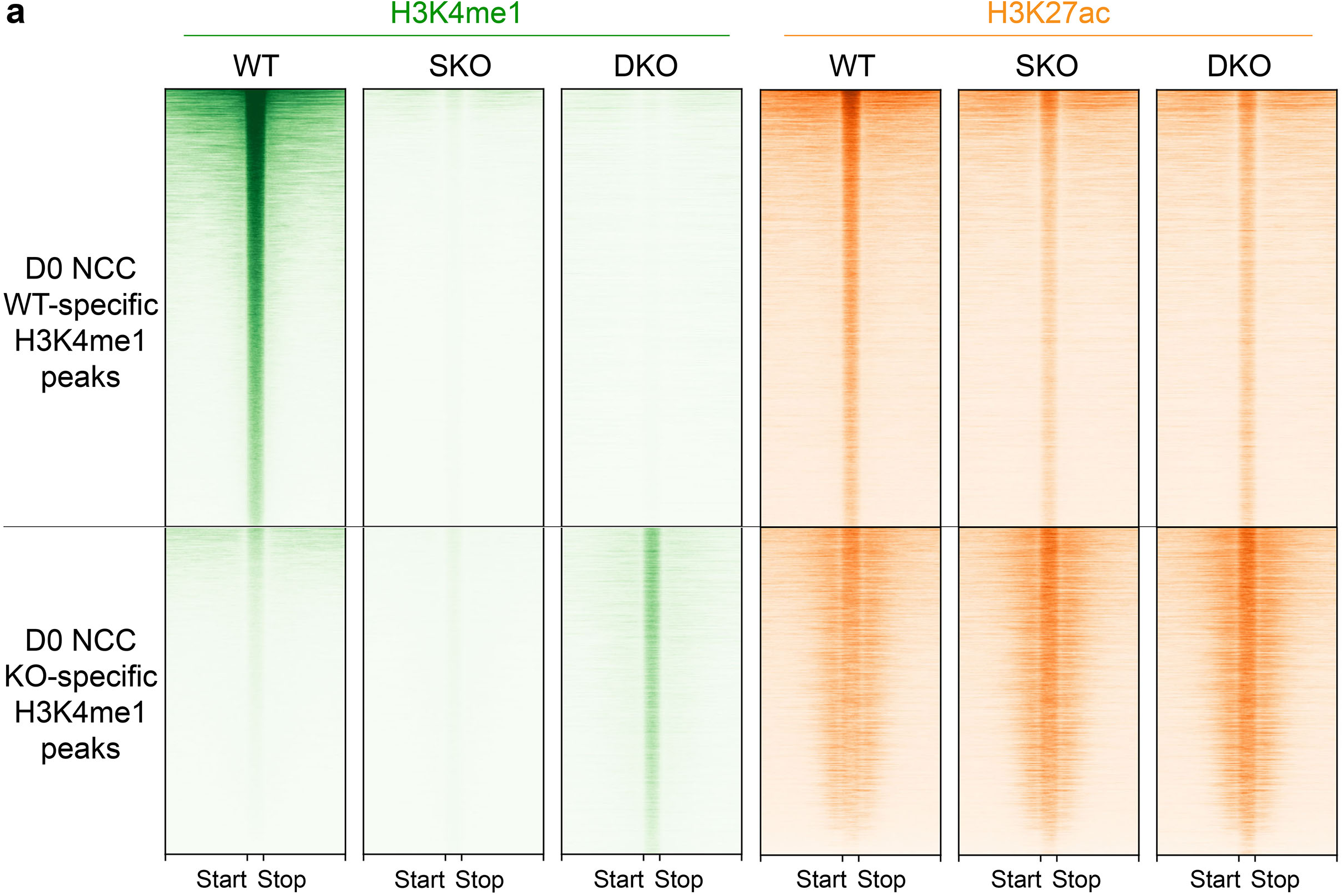
Heatmap of H3K4me1 (green) and H3K27ac (orange) peaks in D0 undifferentiated WT, SKO, and DKO lines, sorted by those that are WT-specific or those that are KO-specific (found only in both SKO and DKO lines).

**Table S1:** Genotype abbreviations for *Col2-Cre* mouse model.

| Genotype | Abbreviation |
| --- | --- |
| *Kmt2c<sup>fl/+</sup> Kmt2d<sup>fl/+</sup> Col2-Cre* | *C<sup>HET</sup> D<sup>HET</sup>* |
| *Kmt2c<sup>fl/fl</sup> Kmt2d<sup>fl/+</sup> Col2-Cre* | *C<sup>KO</sup>* |
| *Kmt2c<sup>fl/+</sup> Kmt2d<sup>fl/fl</sup> Col2-Cre* | *D<sup>KO</sup>* |
| *Kmt2c<sup>fl/fl</sup> Kmt2d<sup>fl/fl</sup> Col2-Cre* | *C<sup>KO</sup> D<sup>KO</sup>* |

**Table S2:**
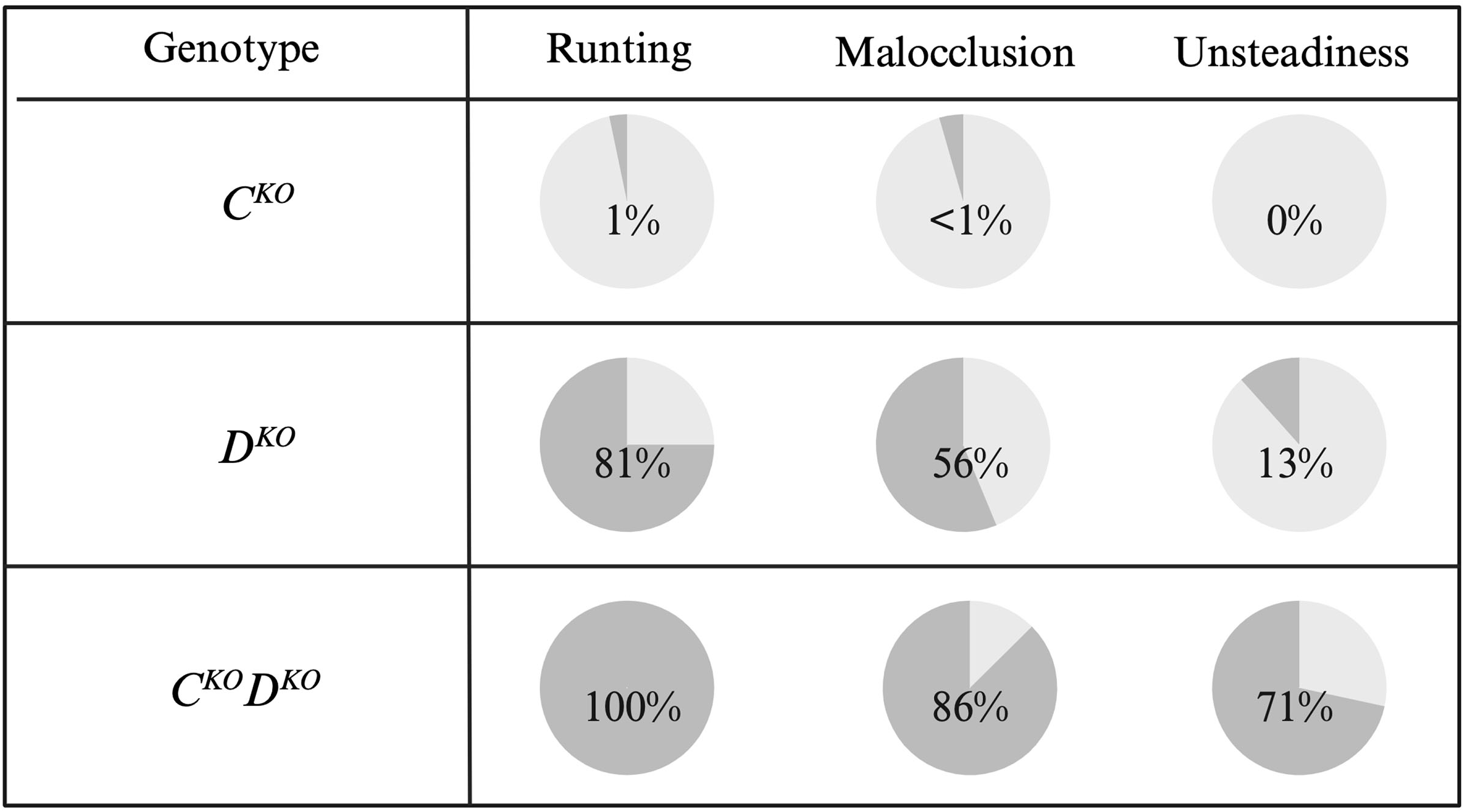
Rates of runting, malocclusion, and unsteadiness observed in weaning age (P24) WT, *C^KO^*, *D^KO^*, and *C^KO^D^KO^* mice (*N=*36, *N=*16, *N=*14, *N=*7).

| Genotype | Runting | Malocclusion | Unsteadiness |
| --- | --- | --- | --- |
| $C^{KO}$ | 1% | <1% | 0% |
| $D^{KO}$ | 81% | 56% | 13% |
| $C^{KO}D^{KO}$ | 100% | 86% | 71% |

**Table S3:**
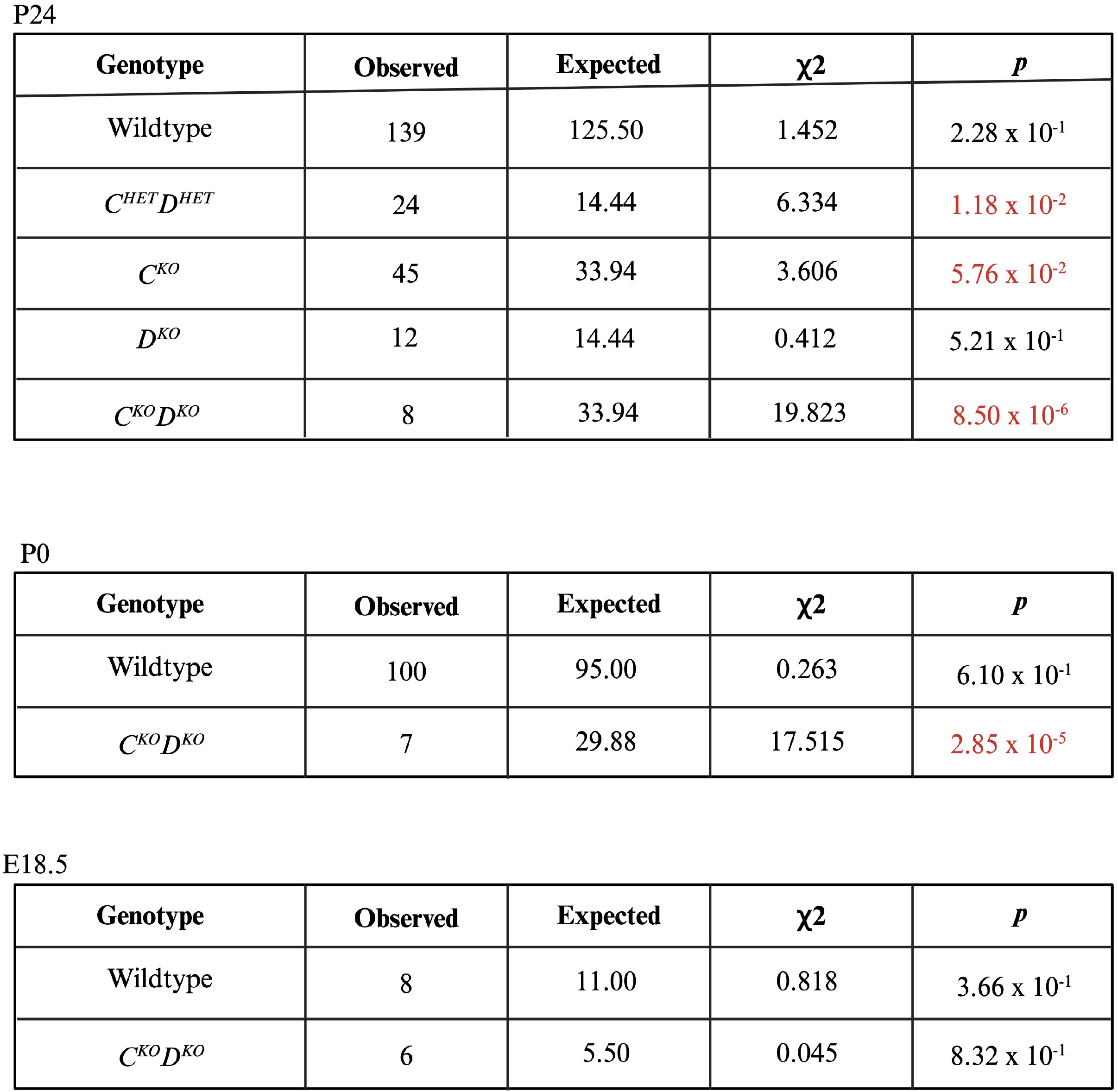
Chi square analysis of all offspring from *Kmt2c (fl/fl) Kmt2d (fl/fl) x Kmt2c (fl/+) Kmt2d (fl/+) Col2-Cre* cross at P24, P0, and E18.5 (*N=*251, *N=*190, *N=*22).

